# Context makes the difference: Temporally Resolved Dopaminergic Teaching Signals Shape Associative Memory in *Drosophila* Larvae

**DOI:** 10.64898/2026.08.17.745210

**Authors:** Denise Weber, Anna-Maria Jürgensen, Juliane Kinnigkeit, Martin P. Nawrot, Andreas S. Thum

## Abstract

Animals can adapt their behavioral responses to environmental cues by learning from experience. This ability relies on the formation and recall of memories that are shaped by beneficial or detrimental consequences and regulated by the dopaminergic system, which is highly conserved across insect species. In the *Drosophila melanogaster* larva, eight of total ∼120 dopaminergic neurons (DANs) innervate the mushroom body (MB), a key center for associative memory. This subset of DANs can be anatomically grouped into two clusters of four cells: the primary protocerebral anterior medial (pPAM) cluster, associated with reward signaling, and the dorsolateral 1 (DL1) cluster, associated with punishment. Such a functional dichotomy is observed in larval and adult *Drosophila* and reflects a fundamental organizational principle of reinforcement learning across invertebrate and even vertebrate species. Aversive reinforcement through high-salt exposure is encoded within the DL1 cluster in a combinatorial and heterogeneous manner, critically involving two neurons, DAN-f1 and DAN-g1. Using temporally precise optogenetic activation and inhibition during olfactory conditioning, we show that these neurons can modulate memory strength and valence. Their effects are most often consistent and predictable, enabling accurate computational modeling of DAN-driven teaching signals. By manipulating the intrinsic physiology of DAN-f1 and DAN-g1 and altering the valence of gustatory input, we are beginning to understand at the single-cell level how dopaminergic activity is systematically adjusted to control memory formation.

## Introduction

Learning is a fundamental cognitive process that enables animals to form experience-dependent memories and predict the consequences of environmental cues and their own actions - capacities essential for survival (Kandel et al 2014, Kandel et al 2012). In behavioral experiments, insects demonstrate a wide spectrum of learning capabilities, ranging from basic associative learning (Cognigni et al 2018, Fiala & Kaun 2024, Giurfa 2015, Hawkins & Byrne 2015, Heisenberg 2003, McGuire et al 2005, Thum & Gerber 2019) to more complex forms such as spatial (Collett et al 2013, Heinze 2017) and social learning (Chittka & Rossi 2022, Giurfa 2012, Leadbeater & Chittka 2007). Their relatively small but highly efficient nervous systems make insects ideal models for studying how experience shapes behavior and for uncovering the underlying neural and molecular mechanisms (Cognigni et al 2018, Menzel 2012, Mizunami et al 2015, Widmann et al 2018).

Among insect models, *Drosophila melanogaster* stands out due to its genetically accessible and well-characterized nervous system (Li et al 2020, Scheffer et al 2020, Schlegel et al 2024, Winding et al 2023). Studies on *Drosophila* larvae, in particular, benefit from the organism’s reduced brain complexity - about 12,000 neurons - and a wealth of robust behavioral assays that allow mechanisms to be studied at single-cell resolution (Apostolopoulou et al 2016, Eschbach et al 2020, Gerber & Hendel 2006, Mancini et al 2019, Pauls et al 2010, Saumweber et al 2018, von Essen et al 2011, Weiglein et al 2019, Winding et al 2023). The availability of powerful genetic tools enables precise manipulations of specific neurons or circuits, while the complete larval brain connectome provides an anatomical ground truth for interpreting functional studies. These features have made the larval *Drosophila* a valuable model for investigating the principles of learning and memory (Eschbach et al 2021, Eschbach et al 2020, Mancini et al 2023, Pauls et al 2010, Saumweber et al 2018, Schleyer et al 2020, von Essen et al 2011, Winding et al 2023). Notably, this system has revealed conserved mechanisms of memory formation across development and species, including compartmentalized reinforcement learning based on dopaminergic neuromodulation (Eschbach et al 2020, Rohwedder et al 2016, Schleyer et al 2020, Weber et al 2025, Weiglein et al 2021).

The current model of MB function for larval and adult *Drosophila* (and insects in general) suggests that its intrinsic Kenyon cells (KCs) integrate sensory input, especially olfactory signals from the antennal lobe but also other modalities, with reinforcement signals to form learned associations (Adel & Griffith 2021, Cognigni et al 2018, Eschbach & Zlatic 2020, Jurgensen et al 2024, Manoim Wolkovitz et al 2026, Modi et al 2020, Parnas et al 2024, Springer & Nawrot 2021, Thum & Gerber 2019, Weber et al 2023b). The KCs transmit sensory information to mushroom body output neurons (MBONs), whose activity is shaped by DANs providing valence-specific teaching signals. In this case these DANs modulate synaptic plasticity at KC-MBON synapses, where aversive learning depresses output from approach-mediating MBONs, and appetitive learning depresses output from avoidance-mediating MBONs. DANs are thus key modulators of learning, updating the value of conditioned stimuli based on reinforcement. In larvae, two small clusters of DANs - each consisting of only four neurons - mediate reward (pPAM) and punishment (DL1) signals, analogous to the related PAM and PPL1 clusters in adults (Rohwedder et al 2016, Saumweber et al 2018, Selcho et al 2009, Weber et al 2025). Under neutral conditions, the baseline activity of both clusters is thought to be balanced, preventing memory retrieval.

Recent model studies (Bennett et al 2021, Gkanias et al 2022, Jurgensen et al 2024, Springer & Nawrot 2021) have emphasized the role of MBON feedback onto the DANs and their role in dynamically computing a prediction error – i.e. the mismatch between expected and actual unconditioned (rewarding or punishing) stimuli -, which can explain fast learning within a single trial as well as the negative acceleration and saturation of learning curves across repeated association trials. MB plasticity is also compartmentalized, facilitating the formation of different types of memories, which are spatially localized within and/or across anatomically distinct compartments of the MB (Aso & Rubin 2016, Bilz et al 2020, Felsenberg et al 2018, Huetteroth et al 2015, Ichinose et al 2015). These compartments interact through synaptic and circuit-level feedback, contributing beyond other functions to the computational flexibility in memory processing (Aso et al 2014, Eichler et al 2017, Eschbach et al 2020, Felsenberg et al 2017, Felsenberg et al 2018, Gkanias et al 2022, Huang et al 2024, Jacob et al 2021, Otto et al 2020, Rachad et al 2025, Saito et al 2026, Springer & Nawrot 2021, Yamada et al 2023, Yamagata et al 2021). Consequently, DANs represent a complex, integrative signaling system. This complexity is also evident in *Drosophila* larvae, where DANs integrate convergent feedback from both aversive and appetitive pathways to compute refined predictive signals to enable future learning (Eichler et al 2017, Eschbach et al 2020).

Given the accessibility and manipulability of DANs, we further investigated their specific roles during different phases of learning in *Drosophila* larvae. Using optogenetics, we selectively activated or inhibited DL1 neurons DAN-f1 and DAN-g1 during the presentation of the conditioned stimulus (CS) and memory retrieval. Our findings show that DANs exhibit baseline activity even without stimulation, and that altering their activity - depending on the timing and pattern - can lead to memory formation in opposite directions. Additionally, we found that optogenetic activation produces a stronger effect than natural stimulation. Finally, we modeled the response dynamics of individual DL1-DANs in a mechanistic computational framework to explore their roles in various mushroom body compartments and their overall contribution to memory formation.

## Results

### Acute optogenetic activation or inactivation of DANs does not impair naïve olfactory and gustatory choice behavior

There are eight DANs in the larval brain that innervate the MB, which can be anatomically classified into two distinct groups: four neurons in the pPAM cluster and four in the DL1 cluster. Previous research has demonstrated a functional division of labor among these larval DANs, with the pPAM cluster primarily mediating dopaminergic reward signals and the DL1 cluster conveying punishment signals (Rohwedder et al 2016, Weber et al 2025). Within each cluster, individual DANs encode distinct yet partially overlapping components of the teaching signal. Specifically, we have identified four DANs within the DL1 cluster - DAN-c1, DAN-d1, DAN-f1, and DAN-g1 - each of which targets distinct compartments of the MB peduncle, lateral appendix, and vertical lobe (Saumweber et al 2018, Weber et al 2025). Notably, DAN-f1/DAN-g1 appear to play a pivotal role in encoding high-salt punishment, as evidenced by their anatomical connectivity, physiological responsiveness to elevated salt concentrations, essentiality for odor high-salt memory, and capacity to functionally substitute for an actual high-salt punishment (Weber et al 2025).

To elucidate in a next step the temporal dynamics encoding the teaching signal in DANs, we employed optogenetic activation and inactivation of DAN-f1 and DAN-g1 under controlled conditions. This approach was chosen based on its technical feasibility and its comparability with our previous work. Therefore, we crossed the split-Gal4 line MB054B for activation experiments with UAS-ChR2XXL, a mutant microbial-type rhodopsin ChR2XXL, which depolarizes neurons artificially (Dawydow et al 2014), and for inhibition approaches with UAS-GtACR2, a light-gated anion-conducting channelrhodopsin from the algae *Guillardia theta* (Mohammad et al 2017). To rule out potential confounding effects on learning and memory, we first examined whether the activity of DAN-f1 and DAN-g1, or the optogenetic blue light stimulation itself, influences innate odor and taste preference (Figure 1 and 2). Optogenetic activation of DAN-f1/DAN-g1 did not affect larval responses to the odors amyl acetate (Figure 1A) and benzaldehyde (Figure 1B), as no differences were observed between blue light-exposed and non-exposed experimental larvae. Similarly, gustatory preference for high-salt (Figure 1C) and fructose (Figure 1D) remained unchanged between these groups. Notably, in two cases, an additional heterozygous control group carrying the UAS-ChR2XXL transgene exhibited a reduction (Figure 1A) or an increase (Figure 1D) in behavior.

**Figure 1:**
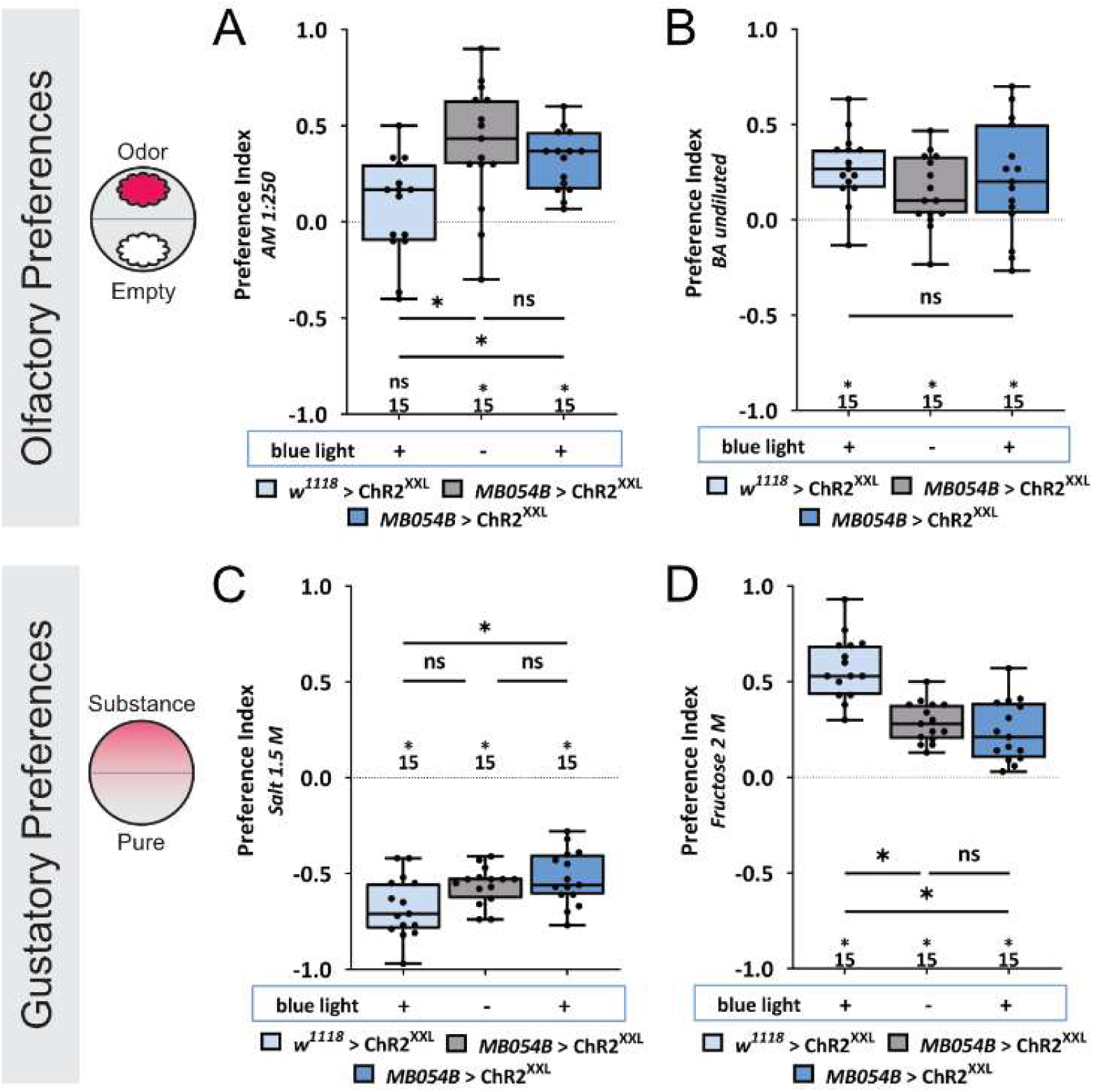
Acute optogenetic activation of DANs does not impair naïve olfactory and gustatory choice behavior. The larval dopaminergic neurons (DANs) can be anatomically categorized into two primary clusters: the primary protocerebral anterior medial (pPAM) and dorsolateral 1 (DL1) clusters, based on the position of their cell bodies. The DL1 cluster comprises four DANs that innervate the c, d, g, and f compartments of the vertical lobe, peduncle, and lateral appendix of the mushroom body (MB). Among these, two DL1 DANs, DAN-f1 and DAN-g1, are included in the expression pattern of the MB054B split-GAL4 driver line. (A, B) To determine whether optogenetic activation of DL1 DAN-f1/DAN-g1 influences naïve odor preference for amyl acetate (AM) and benzaldehyde (BA), we employed the MB054B driver in combination with UAS-ChRXXL. Blue light-induced experimental larvae and uninduced control larvae exhibited comparable behavioral responses to both odors (p > 0.05 for both comparisons). Notably, for amyl acetate, blue-light-exposed UAS-ChRXXL control larvae displayed a distinct response (p < 0.05), showing no odor preference. (C, D) To assess whether optogenetic activation of DL1 DAN-f1/DAN-g1 affects naïve gustatory preference for 1.5 M sodium chloride and 2 M fructose, we again used the MB054B driver in conjunction with UAS-ChRXXL. Blue light-induced experimental larvae and uninduced control larvae demonstrated similar responses to both tastants (p > 0.05 for both comparisons). However, for fructose, blue-light-exposed UAS-ChRXXL control larvae exhibited a stronger preference (p < 0.05). All behavioral data is shown as box-plots. Differences between groups are highlighted by horizontal lines between them. Performance indices different from random distribution are indicated below each box-plot, except for C. Here differences to random distribution are shown above the midline. The sample size of each group (N=15) is given below or above each box-plot. n.s. p > 0.05; * p < 0.05. The source data and results of all statistical tests are documented in Figure 1—source data 1.

**Figure 2:**
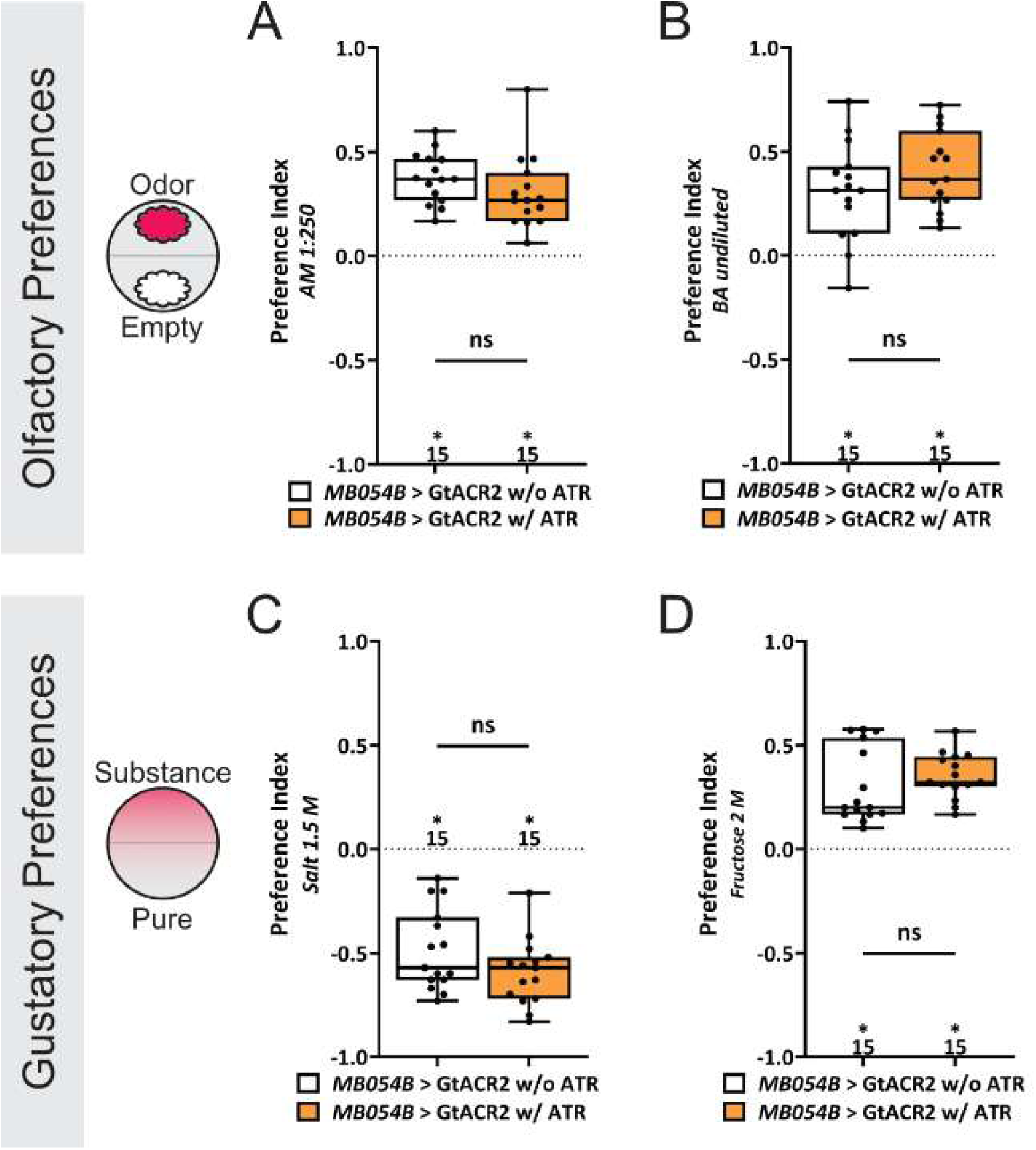
Acute optogenetic inactivation of DANs does not impair naïve olfactory and gustatory choice behavior. (A, B) To investigate whether optogenetic inactivation of the DL1 DAN-f1/DAN-g1 influences naïve odor preference for amyl acetate (AM) and benzaldehyde (BA), we utilized the MB054B driver in combination with UAS-GtACR2. Both, light-induced experimental larvae (fed with all-*trans* retinal, orange) and control larvae (raised without all-*trans* retinal, white) exhibited comparable behavioral responses to both odors (p > 0.05 for both comparisons). (C, D) To assess whether optogenetic inactivation of the DL1 DAN-f1/DAN-g1 affects naïve gustatory preference for 1.5 M sodium chloride and 2 M fructose, we again employed the MB054B driver in combination with UAS-GtACR2. Experimental and control larvae displayed similar responses to both tastants (p > 0.05 for both comparisons). All behavioral data is shown as box-plots. Differences between groups are highlighted by horizontal lines between them. Performance indices different from random distribution are indicated below each box-plot, except for C. Here differences to random distribution are shown above the midline. The sample size of each group (N=15) is given below or above each box-plot. n.s. p > 0.05; * p < 0.05. The source data and results of all statistical tests are documented in Figure 2—source data 1.

A second set of experiments assessed whether inhibition of DAN-f1/DAN-g1 function influences innate odor preference. Suppression of DAN-f1/DAN-g1 activity did not alter responses to amyl acetate (Figure 2A) or benzaldehyde (Figure 2B), as experimental larvae and genetically matched control groups (not exposed to all-*trans*-retinal for GtACR2 activation) displayed comparable odor choice behavior. Likewise, naïve gustatory preference for high-salt (Figure 2C) and fructose (Figure 2D) remained unaffected by DAN-f1/DAN-g1 inhibition. Taken together, these findings show that neither activation nor inhibition of DAN-f1/DAN-g1 alters the innate chemosensory preference of *Drosophila* larvae, suggesting that these neurons do not influence baseline odor or taste choice behavior.

### Optogenetic activation of DL1 DANs is able to overwrite an aversive odor memory

In our previous study, we demonstrated that simultaneous optogenetic activation of DAN-f1/DAN-g1 during odor presentation induces aversive olfactory memory. This memory appears to encode features of natural high-salt stimulation, as evidenced by its retrieval on a high-salt test plate (for a detailed argumentation see (Rahman et al 2026, Schleyer et al 2015, Schleyer et al 2011)). To further investigate the temporal function and dynamics of DAN activation for learning and memory, we analyzed three additional activation paradigms. First, DAN-f1/DAN-g1 activation was paired with the presentation of the odor, which was not associated with high-salt punishment (the unpaired odor CS-). Second, DAN-f1/DAN-g1 activation occurred concurrently with both the odor and high-salt punishment (the paired odor CS+). Third, we extended DAN-f1/DAN-g1 activation throughout the entire training phase, including both odor presentations - the odor paired with high-salt punishment and the odor presented in an unpaired manner (CS+ and CS-).

In the first experimental condition, optogenetic activation of DAN-f1/DAN-g1 during the presentation of the unpaired odor reversed the valence of the memory (Figure 3A). While both control groups exhibited avoidance of the odor associated with high-salt punishment, the experimental group instead avoided the odor paired with optogenetic DAN-f1/DAN-g1 activation (Figure 3A). This indicates that the artificial DAN-f1/DAN-g1-induced memory is capable to overcome the natural high-salt memory. When optogenetic DAN-f1/DAN-g1 activation was paired with the high-salt punishment, no further enhancement of the memory was observed in experimental larvae compared to the control groups. Although a slight trend was present, it did not reach statistical significance (Figure 3B), suggesting that the combination of natural and optogenetically induced memory does not result in a measurable additive effect. Finally, when DAN-f1/DAN-g1 was activated continuously throughout the entire training phase, the aversive memory was abolished (Figure 3C). Experimental larvae exhibited random odor choice behavior during the memory test, unlike the control groups. However, direct statistical comparison among all three groups did not reveal significant differences. Overall, these findings suggest that optogenetically induced DAN-f1/DAN-g1 memory is stronger than naturally acquired high-salt memory and surpasses it under all tested conditions.

**Figure 3.**
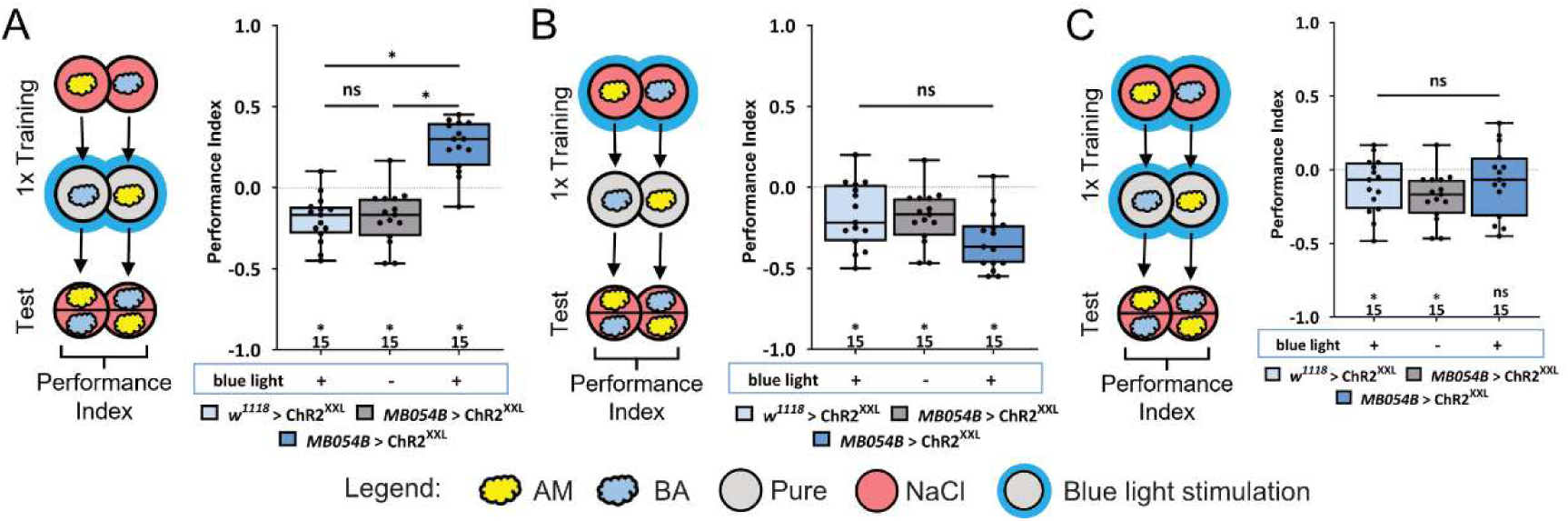
Optogenetic activation of DL1 DANs is able to surpass an aversive odor memory. To investigate whether and how optogenetic activation of DL1 DANs interacts with a high-salt aversive olfactory memory, we employed the MB054B split-GAL4 driver in combination with UAS-ChRXXL. To assess the role of DL1 DAN-f1/DAN-g1 at different training phases, we selectively activated these neurons during (A) the entire presentation of the unpaired odor, (B) the odor paired with high-salt punishment, or (C) both odor presentations, thereby covering the complete training phase. Three experimental groups were analyzed: heterozygous UAS-ChRXXL control larvae exposed to blue light (light blue), MB054B GAL4; UAS-ChRXXL experimental larvae without blue light activation (gray), and MB054B GAL4; UAS-ChRXXL experimental larvae subjected to blue light activation (dark blue). (A) Optogenetic activation during the presentation of the unpaired odor altered memory expression, shifting it from aversive to appetitive for the odor previously associated with high-salt punishment. Blue light-activated experimental larvae exhibited a significantly different response from both control groups (p < 0.05) and were the only group to develop a significant appetitive memory (p < 0.05). (B) Activation of DL1 DAN-f1/DAN-g1 during the presentation of the high-salt-paired odor did not alter aversive memory. All groups exhibited a significant aversive olfactory memory (p < 0.05), with no detectable differences among the three groups (p > 0.05). (C) Activation of DL1 DAN-f1/DAN-g1 throughout the entire training phase impaired the formation of a high-salt aversive olfactory memory (p > 0.05). However, no significant differences in memory performance were detected between the three groups (p > 0.05). All behavioral data is shown as box-plots. Differences between groups are highlighted by horizontal lines between them. Performance indices different from random distribution are indicated below each box-plot. The sample size of each group (N=15) is given below each box-plot. n.s. p > 0.05; * p < 0.05. The source data and results of all statistical tests are documented in Figure 3—source data 1.

### Optogenetic activation of DL1 DANs is able to write a context independent aversive odor memory

Several studies showed that in the standard aversive olfactory learning paradigm larvae not only remember the value of the reinforcement, but also its quality (Rahman et al 2026, Schleyer et al 2015, Schleyer et al 2011). The findings show that behavior is best conceptualized as a memory-driven search for a more favorable condition. High-salt memory is only expressed in the presence of high-salt; when tested on a pure agarose plate, lower salt concentrations or even a bitter quinine-containing plate, the learned behavior is not shown. This indicates that the learned response is based on a relative assessment, where larvae recall both the intensity and identity of the punishment experienced during training and compare this stored representation with the current test conditions. These findings provide a methodological framework for testing the specific identity of the larval optogenetically written aversive memory by varying test conditions. Following this rationale, we examined the content of the optogenetically induced DAN-f1/DAN-g1 memory by testing larval recall on plates containing high-salt, quinine, fructose, or pure agarose (Figure 4A–D). Unexpectedly, in all four conditions, experimental larvae exhibited robust aversive olfactory memory, which differed significantly from both control groups. This suggests that, unlike naturally acquired high-salt memory, the optogenetically induced DAN-f1/DAN-g1 memory is independent of the test context and does not encode a specific stimulus quality. Alternatively, it may even encode multiple qualities simultaneously, allowing for recall across diverse test conditions.

**Figure 4:**
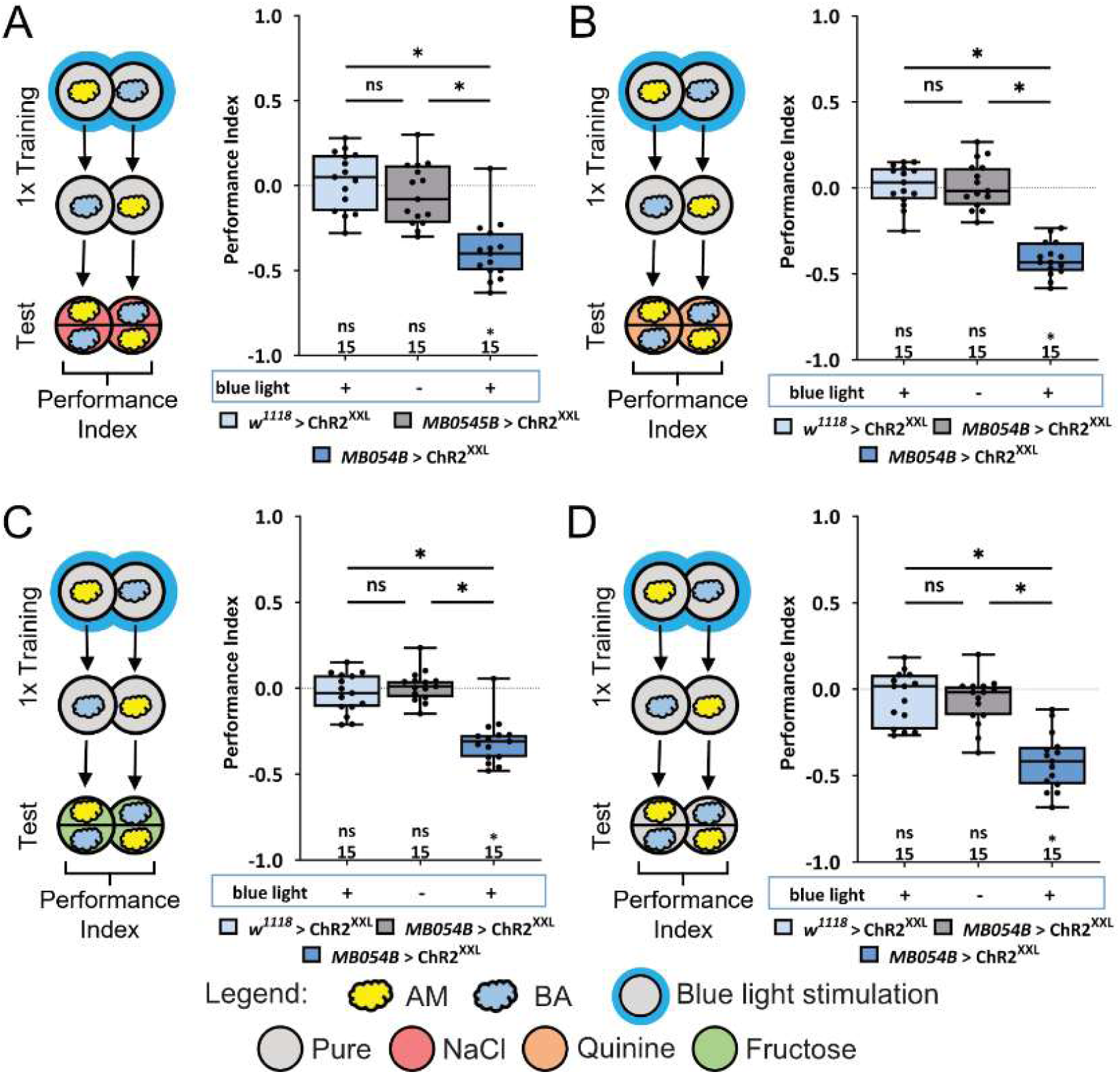
Optogenetic activation of DL1 DANs is able to write a context independent aversive odor memory. To investigate whether the memory induced by optogenetic activation of DL1 DANs encodes qualitative aspects of punishment, we conducted four independent experiments. Experimental larvae were tested on substrates containing either high-salt (A), bitter quinine (B), fructose (C), or pure agarose (D). Three groups were compared: heterozygous UAS-ChRXXL control larvae exposed to blue light (light blue), MB054B split-GAL4; UAS-ChRXXL experimental larvae without blue light activation (gray), and MB054B split-GAL4; UAS-ChRXXL experimental larvae that received blue light activation (dark blue). (A–D) Our results demonstrate that the memory induced by DL1 DAN-f1/DAN-g1 activation was recalled independently of the test substrate. In all conditions, only the experimental group exhibited a significant aversive olfactory memory (p < 0.05), which was consistently different from both control groups (p < 0.05), while the control groups distributed randomly (p > 0.05). These findings suggest that under the tested conditions, optogenetic activation of DL1 DAN-f1/DAN-g1 is capable of inducing a memory that can be retrieved regardless of the test situation, indicating its context-independent nature. All behavioral data is shown as box-plots. Differences between groups are highlighted by horizontal lines between them. Performance indices different from random distribution are indicated below or above each box-plot. The sample size of each group (N=15) is given for each box-plot. n.s. p > 0.05; * p < 0.05. The source data and results of all statistical tests are documented in Figure 4—source data 1.

### DAN inhibition during training exerts phase-dependent and opposing effects on odor memory

In our previous study, we demonstrated that acute optogenetic inhibition of DAN-f1 and DAN-g1 throughout the entire training phase impairs odor high-salt memory (Weber et al 2025). To further elucidate the temporal dynamics and functional role of DAN inactivation, we implemented three additional inhibition paradigms alongside a replication of the previously published experiment (Figure 5).

**Figure 5:**
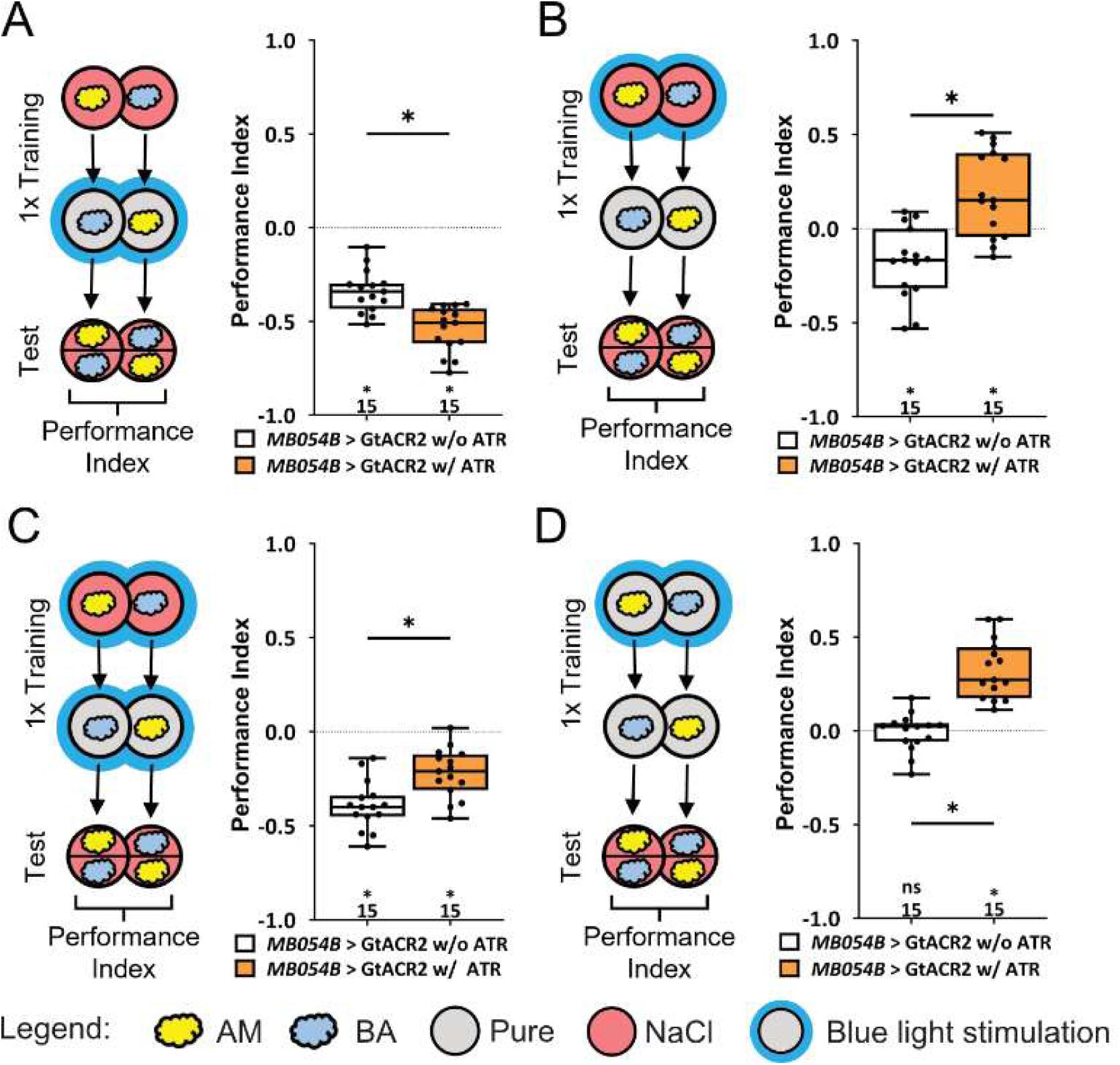
Context dependent inhibition of DL1 DAN activity changes aversive odor memories in opposing ways. To investigate whether optogenetic inactivation of DL1 DANs influences high-salt aversive olfactory memory, we utilized the MB054B split-GAL4 driver in combination with UAS-GtACR2. To examine the role of DAN-f1/DAN-g1 at distinct training time points, we selectively inhibited these neurons during (A) the entire presentation of the unpaired odor, (B) the odor paired with high-salt punishment, (C) both odor presentations, encompassing the entire training phase, or (D) the presentation of the first odor in a modified protocol that utilized pure agarose plates, omitting the high-salt teaching signal. Two experimental groups were tested: MB054B split-GAL4; UAS-GtACR2 control larvae that did not receive all-*trans* retinal in their diet (white) and MB054B split-GAL4; UAS-GtACR2 experimental larvae that were fed all-*trans* retinal throughout development (orange). (A) Optogenetic inactivation during the presentation of the unpaired odor enhanced aversive memory, leading to stronger avoidance of the odor associated with high-salt punishment (p < 0.05). (B) Inactivation of DAN-f1/DAN-g1 during the presentation of the high-salt-paired odor significantly weakened aversive olfactory memory, even reversing it into an appetitive memory (p < 0.05). (C) Inhibition of DAN-f1/DAN-g1 throughout the entire training phase reduced high-salt aversive olfactory memory compared to the control group (p < 0.05). However, a residual but significantly detectable aversive memory remained (p < 0.05). (D) Interestingly, even inhibiting DAN-f1/DAN-g1 function in the absence of any teaching stimulus influenced larval learning. Optogenetic DAN inhibition paired with an odor alone was sufficient to induce an appetitive olfactory memory (p < 0.05), which, in contrast to the control group, was significantly different from a random distribution (p < 0.05). All behavioral data is shown as box-plots. Differences between groups are highlighted by horizontal lines between them. Performance indices different from random distribution are indicated below or above each box-plot. The sample size of each group (N=15) is given for each box-plot. n.s. p > 0.05; * p < 0.05. The source data and results of all statistical tests are documented in Figure 5—source data 1.

First, DAN-f1/DAN-g1 inhibition was applied during the presentation of the odor that was not paired with high-salt punishment (the unpaired odor, CS-). Second, inhibition was synchronized with the presentation of the odor paired with high-salt punishment (paired odor, CS+). Third, DAN-f1/DAN-g1 inhibition was extended to the entire training phase, encompassing both odor presentations, to reproduce the previously reported findings. Finally, we tested a condition in which high-salt punishment during training was fully replaced with a pure agarose plate combined with simultaneous optogenetic inhibition of DAN-f1/g1.

In the first experimental condition, optogenetic inhibition of DAN-f1/DAN-g1 during the presentation of the unpaired odor significantly enhanced memory strength (Figure 5A). Control larvae with the same genotype but without all*-trans* retinal feeding avoided the odor associated with high-salt punishment. However, this avoidance response was more pronounced in the all*-trans* retinal fed larvae (Figure 5A). When optogenetic inhibition of DAN-f1/DAN-g1 was paired with high-salt punishment, a reduction in memory strength was observed in experimental larvae compared to the control group (Figure 5B). Continuous inhibition of DAN-f1/DAN-g1 throughout the entire training phase produced a similar effect to that previously reported, leading to a diminished aversive odor high-salt memory (Figure 5C). However, in contrast to the experimental group in the previous experiment, the larvae still exhibited a reduced aversive memory rather than a slightly positive memory (Figure 5B and C). Finally, the fourth experiment demonstrated that optogenetic inhibition of DAN-f1/DAN-g1 alone was sufficient to induce a positive memory (Figure 5D). Experimental larvae, unlike the control group that distributed randomly, actively approached the odor associated with DAN inhibition, indicating a positive association. Overall, these results suggest that DAN-f1/DAN-g1 neurons exhibit a basal level of activity. Blocking this activity appears to have a rewarding effect on the larvae (Figure 5A and D) or reduces the effectiveness of a simultaneously presented high-salt punishment (Figure 5B and C).

### Acute manipulation of DL1 DANs affects the recall of odor high-salt memory

In the previously described experiments, we investigated the impact of DAN-f1/DAN-g1 manipulation during the training phase. In the next step, we examined its role specifically during the test phase, focusing on memory recall (Figure 6 and 7).

**Figure 6.**
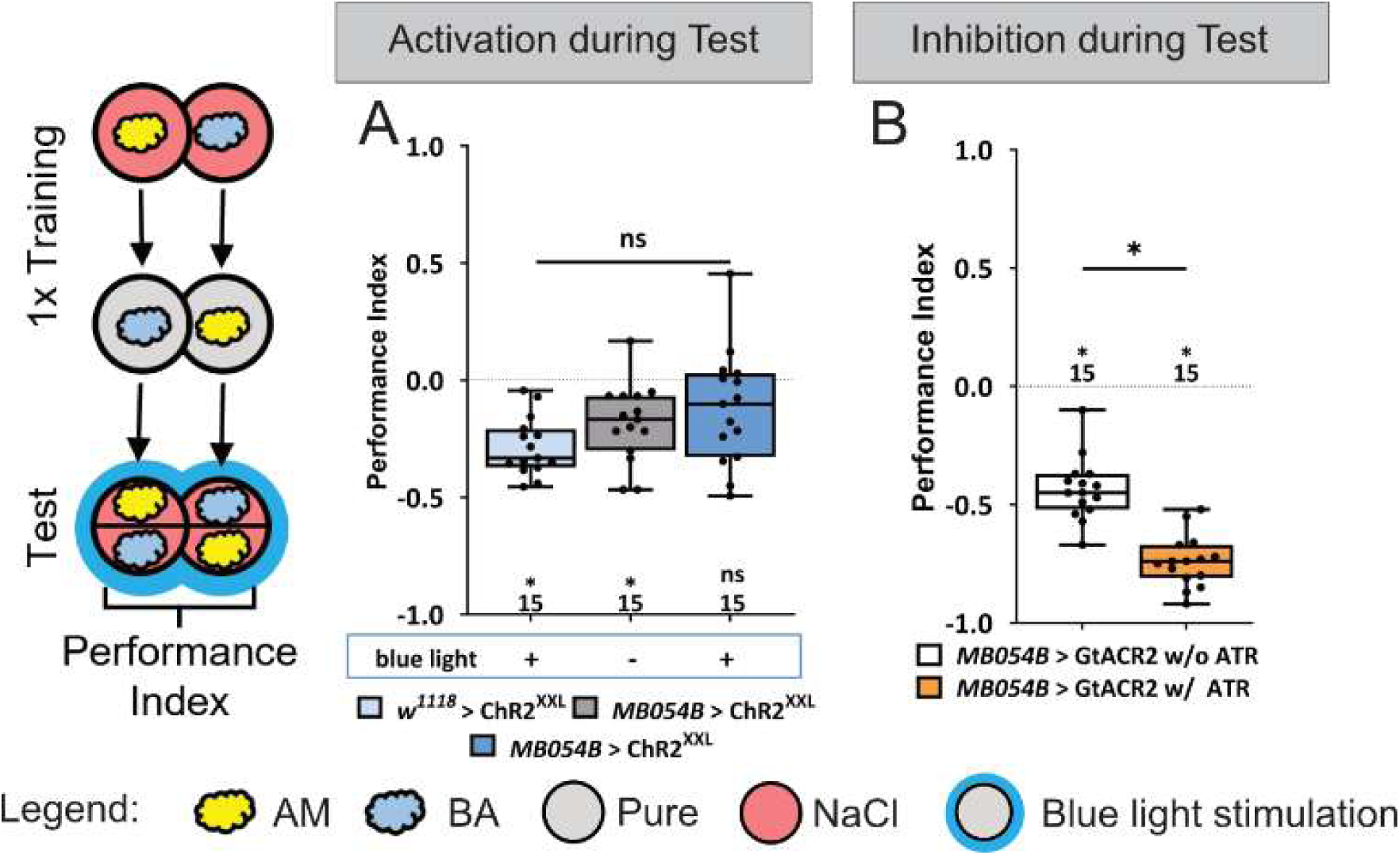
Acute manipulation of DL1 DANs also affects the recall of odor high-salt memory. To determine whether optogenetic manipulation of DL1 DANs during the test phase influences high-salt aversive olfactory memory retrieval, we employed the MB054B split-GAL4 driver in combination with either UAS-ChRXXL for activation (A) or UAS-GtACR2 for inactivation (B). Our findings revealed opposing effects on memory expression. (A) Optogenetic activation of DAN-f1/DAN-g1 during the test phase abolished aversive memory (p > 0.05) compared to the two control groups. While both control groups exhibited significant aversive memory (p < 0.05), no significant differences were detected among the three groups (p > 0.05). (B) In contrast, optogenetic inactivation of DAN-f1/DAN-g1 during the test phase significantly enhanced high-salt aversive olfactory memory (p < 0.05). These results suggest that DAN-f1/DAN-g1 not only contribute to encoding a teaching signal during learning but also play a role in memory retrieval. For activation experiments (A), three experimental groups were analyzed: heterozygous UAS-ChRXXL control larvae exposed to blue light (light blue), MB054B split-GAL4; UAS-ChRXXL experimental larvae without blue light activation (gray), and MB054B split-GAL4; UAS-ChRXXL experimental larvae subjected to blue light activation (dark blue). For inactivation experiments (B), two groups were tested: MB054B split-GAL4; UAS-GtACR2 control larvae that did not receive all-*trans* retinal in their diet (white) and MB054B split-GAL4; UAS-GtACR2 experimental larvae that were fed all-*trans* retinal throughout development (orange). All behavioral data is shown as box-plots. Differences between groups are highlighted by horizontal lines between them. Performance indices different from random distribution are indicated below (in A) or above (in B) each box-plot. The sample size of each group (N=15) is given for each box-plot. n.s. p > 0.05; * p < 0.05. The source data and results of all statistical tests are documented in Figure 6—source data 1.

**Figure 7:**
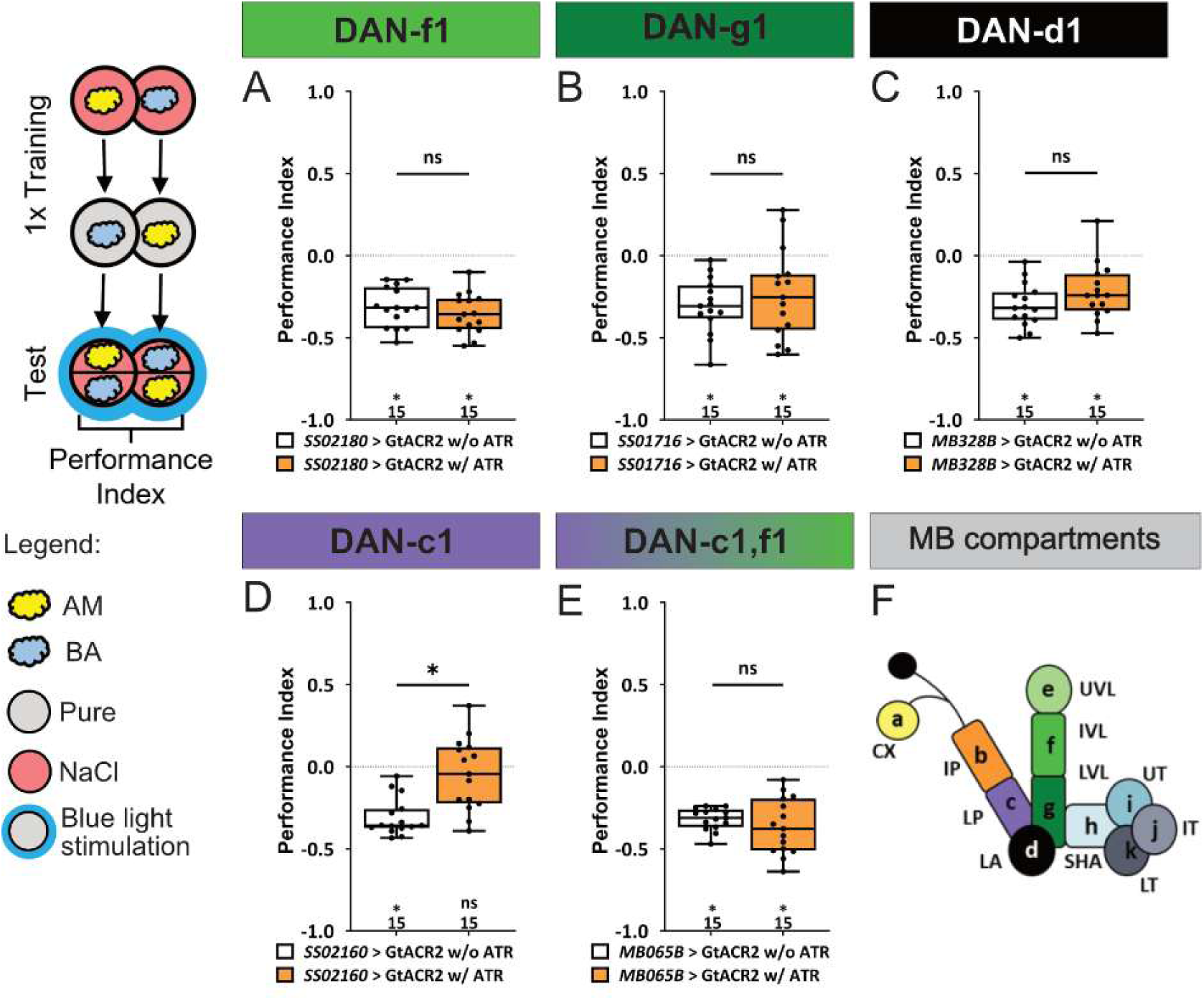
Acute manipulation of individual DL1 DANs affects the recall of odor high-salt memory. To determine whether optogenetic inactivation of individual DL1 DANs specifically during the test phase influences high-salt aversive olfactory memory retrieval, we used SS02180 (DAN-f1), SS01716 (DAN-g1), MB328B (DAN-d1), SS02160 (DAN-c1), and MB065B (combination of DAN-c1 and DAN-f1) split-GAL4 drivers together with UAS-GtACR2. For all experiments two groups were tested: split-GAL4; UAS-GtACR2 control larvae that did not receive all-*trans* retinal in their diet (white) and split-GAL4; UAS-GtACR2 experimental larvae that were fed all-*trans* retinal throughout development (orange). (A-C, E) Inactivation of DAN-f1, DAN-g1, DAN-d1, or the combination of DAN-c1/DAN-f1 during test did not reduce the aversive olfactory high-salt memory (p < 0.05). All experimental groups performed as their respective control group (p > 0.05). (D) Inactivation of DAN-c1 alone impaired aversive olfactory high-salt memory (p > 0.05). The experimental larvae behaved differently compared to the control group (p < 0.05). (F) The larval MB is organized into 11 compartments: CX calyx; IP and LP intermediate and lower peduncle; LA lateral appendix; UVL, IVL, and LVL upper, intermediate, and lower vertical lobe; SHA, UT, IT, LT shaft as well as upper, intermediate, and lower toe of the medial lobe. Single-letter synonyms of compartment names are given as “a–k”. These letters are used to indicate compartment innervation by the MB input and output (Saumweber et al 2018). All behavioral data is shown as box-plots. Differences between groups are highlighted by horizontal lines between them. Performance indices different from random distribution are indicated below each box-plot. The sample size of each group (N=15) is given below each box-plot. n.s. p > 0.05; * p < 0.05. The source data and results of all statistical tests are documented in Figure 7—source data 1.

Optogenetic activation of DAN-f1/DAN-g1 during the test phase in the odor high-salt paradigm led to a random distribution of experimental larvae, in contrast to both control groups, indicating a loss of memory expression (Figure 6A). However, the performance indices of the three groups did not differ significantly. Conversely, optogenetic inhibition of DAN-f1/DAN-g1 during the test phase enhanced memory recall (Figure 6B). In this condition, experimental larvae exhibited stronger avoidance of the odor associated with high-salt punishment compared to control groups, which also displayed aversive memory. These findings suggest that DAN manipulation affects not only the training phase but also the test phase, indicating that DANs in larvae do more than simply mediate a teaching signal, as previously assumed.

To further investigate the role of DL1 cluster DANs during memory recall, we conducted a detailed analysis of each of the four individual DL1 neurons (Figure 7). Specifically, we inhibited DAN-f1, DAN-g1, DAN-d1, DAN-c1, or the combination of DAN-c1/DAN-f1 during the test phase using UAS-GtACR2. A schematic representation of MB compartment innervation by these DANs is provided in Figure 7F. The results indicate that inhibition of individual DANs generally had little effect on aversive memory expression (Figure 7A–C and E). However, one exception was observed: inhibition of DAN-c1 led to a significant behavioral difference between experimental and control larvae (Figure 7D). Interestingly, this effect was the opposite of the previously observed DAN-f1/DAN-g1 inhibition (Figure 6B), as it resulted in complete memory impairment rather than enhancement. The effect appears to be specific to memory, as DAN-c1 inhibition does not induce any detectable deficits in sensory acuity for the odors amyl acetate and benzaldehyde or the tested high-salt concentration (Figure 7-figure supplement 1)). In summary, these findings suggest that the activity of DL1 DAN neurons plays a functional role in odor high-salt memory not only during learning but also during the test phase.

### A circuit model links time-resolved DAN activity to compartment-specific mushroom body plasticity

The behavioral experiments revealed that manipulation of DAN-f1/g1 does not simply scale aversive learning, but can also overwrite, weaken, or even reverse memory valence depending on the training context. These effects are difficult to interpret from the activity of individual DANs alone, because each DAN acts within a compartmental MB circuit in which parallel KC–MBON plasticity and feedback interactions may jointly determine behavioral output. We therefore implemented a circuit model of the larval MB to test whether experimentally observed individual DAN response dynamics are sufficient to account for our behavioral effects of high-salt punishment, optogenetic DAN activation, and DAN silencing.

As a first step, we independently fitted the parameters of a simple dynamical neuron model for each of the four DL1 DANs and for the pPAM cluster to the previously measured calcium responses evoked by 5-10s high-salt or fructose stimulation (Figure 8) (Weber et al 2025). The fitted responses captured well the distinct temporal response, adaptation profiles, and sensitivities of individual DL1 DANs and the pPAM cluster DANs, including the strong salt responses of DAN-c1 and DAN-d1, the weaker salt responses of DAN-f1 and DAN-g1, and the opposing high-salt and fructose sensitivity of the pPAM cluster.

**Figure 8:**
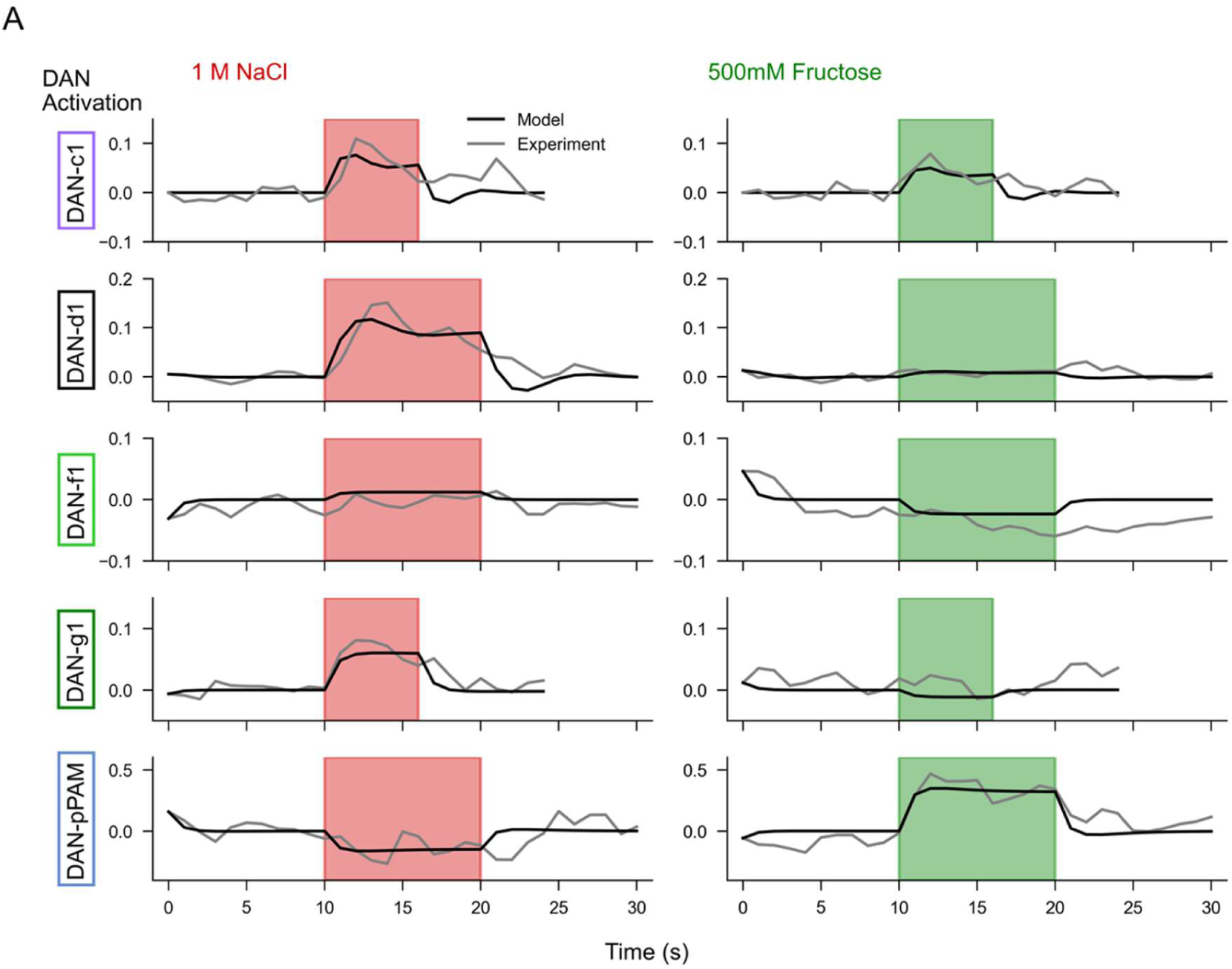
Time-varying DAN calcium traces in responses to stimulation with gustatory high-salt or fructose in experiment and model. A unique neuron model was implemented for each individual DL1 (DAN-c1, DAN-d1, DAN-f1, DAN-g1), and one neuron model to represent the pPAM cluster. Calcium recordings of the individual DL1 neurons and of the collective pPAM cluster allowed to quantify their responses to gustatory high-salt (1 M) or fructose (500 mM) in the intact animal (Weber et al., 2025). The neuron model parameters - time constant of the neuronal responses to the stimulation, the weight of the sensory input *winp_DAN* (salt or fructose), and the magnitude and time constant of neuronal adaptation - were fitted by minimizing the difference between the model response (black) and experimental data (gray). Red/green shaded area denotes the salt/fructose stimulation, respectively. Experimental data from (Weber et al 2025).

The basic circuit model in Figure 9A represents the larval MB as parallel compartments in which odor-evoked KC activity converges onto MBONs through plastic KC–MBON synapses. Using this framework, we simulated the relevant training conditions by activating DANs with high-salt, optogenetically activating DAN-f1/g1, or optogenetically silencing DAN-f1/g1 during odor presentation (Figure 9B, C). The experimentally constrained DAN response models (Figure 8) allowed for a biologically realistic simulation of compartment-specific teaching signals (Figure 9B). Coincident activity of KCs and a single DAN induced depression of the corresponding KC–MBON synapses (Figure 9B), reflecting the compartmental organization of dopamine-modulated plasticity in the MB (Davidson et al 2023, Eschbach et al 2020, Hige et al 2015, Saumweber et al 2018). In this architecture, DL1-DAN-driven depression reduces the activity of approach-promoting MBON channels and thereby favors avoidance, whereas pPAM-driven depression reduces avoidance-promoting MBON output and thereby favors approach. The model output, as a proxy to larval behavior, is the avoidance bias, computed from the relative activity of avoidance-promoting and approach-promoting MBONs. Positive values indicate a bias toward odor avoidance behavior, whereas negative values indicate a bias toward odor approach behavior. This model provides a mechanistic bridge between experimentally measured DAN response dynamics, compartment-specific synaptic plasticity, and the learned behavior.

**Figure 9:**
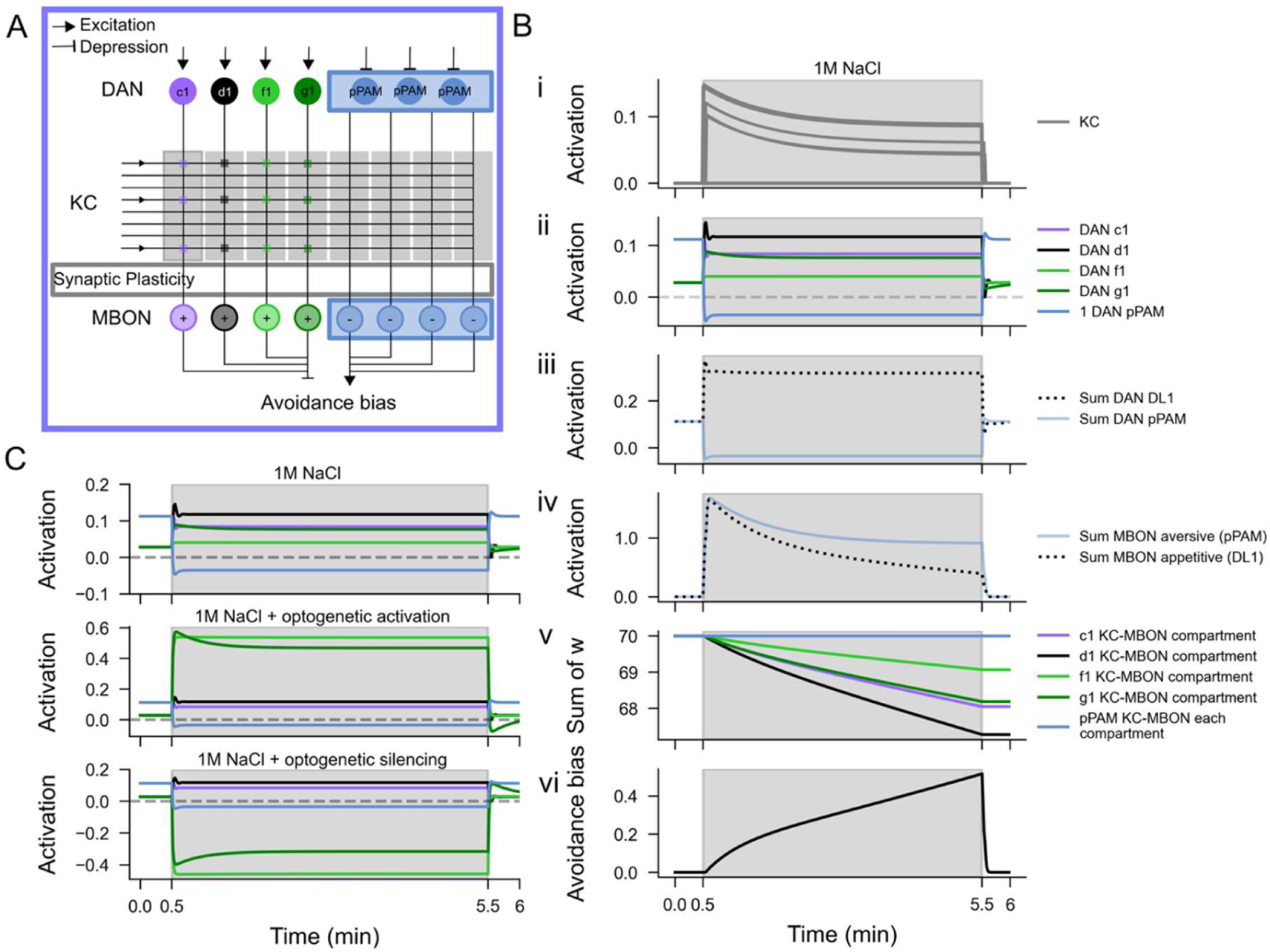
MB circuit model shows DAN-mediated compartment specific synaptic plasticity and the resulting avoidance bias. (A) Basic circuit model. Plasticity at KC-MBON synapses is implemented as a learning rule that reduces synaptic weights when odor-evoked KC activation coincides with DAN activity within a specific compartment (DAN-c1, DAN-d1, DAN-f1, DAN-g1, pPAM). This results in an imbalance in the activity of MBONs associated with approach- and avoidance-promoting compartments. Each DL1 or pPAM DAN innervates a distinct compartment and modulates plasticity locally. (B) Synaptic plasticity in each compartment as expressed in the dynamic reduction of synaptic strength (v) is driven by the coincidence of KC (i) and compartment-specific DAN (ii,iii) activity, leading to skewed MBON output (iv). This imbalance alters the relative strength of approach- and avoidance-promoting MB output to downstream circuits, controlling aversive behavior as expressed in the increasing avoidance bias (vi). (C) In the model, and depending on the experimental protocol simulated, we compare DAN activation either by high-salt stimulation, by means of optogenetic activation (DAN-f1/DAN-g1), or in optogenetic silencing (DAN-f1/DAN-g1).

### Cross-compartmental feedback is required to explain valence reversal during punishment learning

We next asked whether independently operating MB compartments are sufficient to explain the behavioral effects observed after optogenetic manipulation of DAN-f1/g1 during learning. The model was evaluated against a set of qualitative behavioral constraints. Odor presentation alone should not induce a strong learned bias. Pairing odor with high-salt should induce avoidance. Pairing odor with optogenetic activation of DAN-f1/g1 should induce strong avoidance, consistent with the experimentally observed artificial aversive memory (Figure 3). Conversely, pairing odor with optogenetic silencing of DAN-f1/g1 should shift memory valence toward approach (Figure 5). Finally, the model should account for the most informative condition, in which high-salt is paired with simultaneous silencing of DAN-f1/g1 and the behavioral output shifts away from normal aversive memory (valence reversal, Figure 5).

The basic model (Model 1, Figure 9A) reproduced the straightforward aversive-learning conditions as expressed in the avoidance bias (Figure 10A), including avoidance after high-salt training and after optogenetic DAN-f1/g1 activation. However, it failed to reproduce the valence reversal observed when DAN-f1/g1 were silenced during punishment learning (Figure 10A). This failure is informative because it shows that the behavioral reversal cannot be explained by independent compartmental plasticity driven only by the fitted DAN responses. In such a purely parallel model, silencing part of the DL1 teaching signal can reduce aversive learning, but it does not actively generate the reward-like signal required to reverse valence. We therefore tested whether adding novel compartmental interactions to our model based on the larval connectome (Eichler et al 2017, Eschbach et al 2020) could account for this discrepancy. Removing salt-driven activation of DAN-f1 (Figure 10-figure supplement 1A, Model 2) had little effect, consistent with the weak fitted salt response of this neuron. Adding inhibitory interactions between approach- and avoidance-promoting MBON channels changed the strength of the predicted avoidance bias but did not resolve the reversal condition (Figure 10-figure supplement 1B, Model 3). Similarly, a more generic MBON-to-DAN feedback motif was insufficient to reproduce the full behavioral pattern (Figure 10-figure supplement 1C, Model 4). Thus, neither altered DAN-f1 salt sensitivity nor nonspecific interactions between MBON channels were sufficient to explain the experimental observations.

**Figure 10:**
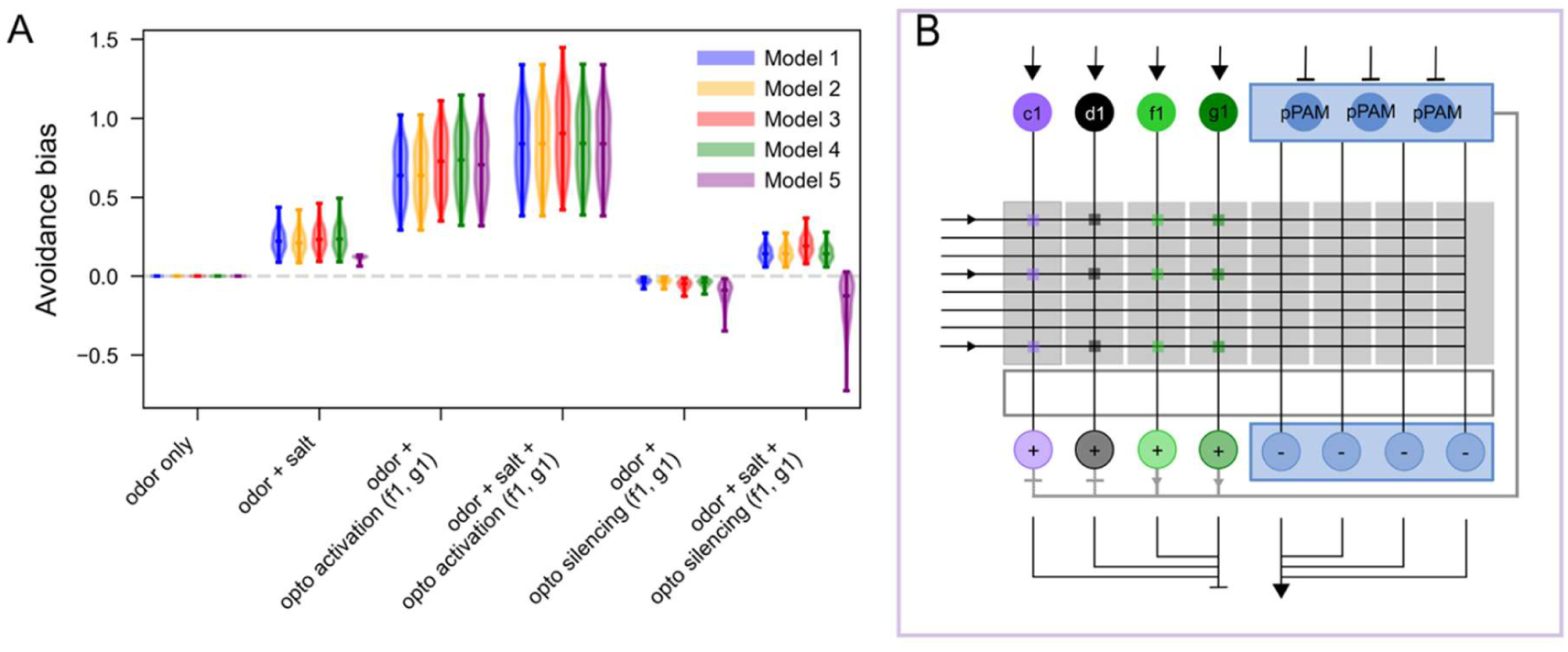
Simulated olfactory learning experiments extended circuit models. The performance of the basic model (Model 1, Figure 9A) is compared to the performance of four alternative circuits in different learning experiments where an odor is paired with stimulation, activation or inactivation of specific DANs. (A) Comparison of avoidance bias across five different models in six different simulated experiments where an odor is presented alone (odor only) or is paired with high-salt, optogenetic activation (opto activation) or optogenetic silencing (opto silencing) of the DAN-f1/g1, or with combination high-salt and silencing. Violin plots show the distribution of the avoidance bias across 200 model instances with the median plotted as line and the minimum and maximum data points plotted as whiskers. (B) Circuit sketch of Model 5 that could qualitatively reproduce the experimental results across all experimental protocols, including the valence reversal (Figure 5) when pairing odor with high-salt while silencing DAN-f1 and DAN-g1.

Only a circuit variant with specific cross-compartmental feedback reproduced the valence reversal (Figure 10-figure supplement 1D, Model 5). In this model, MBON output from the f1/g1 compartments provides excitatory feedback onto pPAM DANs, whereas MBON output from the c1/d1 compartments provides balancing inhibitory input. Under normal high-salt training, this circuit remains biased toward aversive learning. However, when DAN-f1/g1 are silenced during punishment learning, the balance of the circuit is shifted such that pPAM activity can be recruited, producing an approach-promoting teaching signal and reversing the predicted behavioral valence (Figure 10B). The model comparison therefore suggests that valence reversal is not simply caused by the absence of DAN-f1/g1-mediated punishment signaling. Instead, it predicts a circuit-level rebalancing mechanism in which perturbation of aversive DL1 compartments can recruit reward-like pPAM teaching signals through cross-compartmental MBON–DAN feedback. This provides a testable prediction: the f1/g1-associated MBON pathway should be able to excite pPAM DANs directly or indirectly, while c1/d1-associated pathways should counterbalance this excitation in the intact circuit. More generally, the simulations indicate that the behavioral effects of DAN-f1/g1 manipulation emerge from the interaction between measured DAN response dynamics and recurrent MB circuit architecture, rather than from the activity of individual DANs alone.

## Discussion

The dopaminergic system is an evolutionarily conserved signaling pathway that plays a central role in associative learning across both invertebrate and vertebrate species. Our study shows that temporally structured dopaminergic activity is a key determinant of how aversive olfactory memories are written, expressed, and even reversed in *Drosophila* larvae using temporally optogenetic activation and inhibition of DL cluster DAN-f1/DAN-g1. Our findings, considered alongside published data, suggest that that individual larval DANs provide distinct, temporally resolved teaching signals that interact across MB compartments to generate flexible, valence-reversible memories. Furthermore, these findings reveal fundamental organizational principles of DAN–mediated teaching signals that likely extend beyond larvae and insects.

### Optogenetically written memory is context-independent

A remarkable observation was that the aversive memory written via the optogenetic DAN-f1/DAN-g1 activation was expressed across all test conditions including high-salt, quinine, fructose, and plain agarose (Figure 4). In contrast, aversive memory written via a real-world physical high-salt stimulus is only retrieved when test conditions match the punishing context (Rahman et al 2026, Schleyer et al 2015, Schleyer et al 2011). Larvae therefore appear to encode not only the learned valence of a cue (good or bad), but also the qualitative identity of the reinforcing stimulus (e.g. high-salt) in the established memory trace (Schleyer et al 2015). Consistent with this view, aversive quinine memories are retrieved only when quinine is present during the test, whereas appetitive memories induced by fructose or aspartic acid are selectively blocked only by the corresponding reinforcer itself (Rohwedder et al 2016, Schleyer et al 2015, Schleyer et al 2011). In contrast optogenetic DAN-f1/DAN-g1 activation seems to write a generic aversive engram. One possibility is that artificial DAN activation, unlike natural reinforcement, produces stronger KC–MBON depression that bypasses or overrides the quality-dependent gating of memory expression. This interpretation is supported by work in the adult MB, where pairing odor presentation with optogenetic activation of the PPL1-γ1pedc DAN induces robust, stimulus-specific long-term depression at the corresponding KC-MBON synapses (Hige et al 2015). Alternatively, artificial activation may engage broader KC-MBON depression across MB compartments f and g, thereby generating a mixed-content or generalized memory. Such an interpretation is consistent with studies in adult *Drosophila* showing that distinct MB compartments contribute to memories associated with sweet taste, water, and the nutrient value of sugar (Burke et al 2012, Huetteroth et al 2015, Ichinose et al 2015, Lin et al 2014, Liu et al 2012). Moreover, optogenetic activation of individual DANs has revealed that different adult MB compartments support memories with “content”, such as distinct strengths, acquisition kinetics, and durations (Aso & Rubin 2016). In line with this view, artificial co-activation of nearly all reward-encoding adult MB compartments can induce an appetitive memory that substantially exceeds the strength of naturally induced sugar memory, as demonstrated by its persistence even after seven days of ad libitum feeding following training (Huetteroth et al 2015). Finally, MB downstream or feedback circuits may be differentially affected by the magnitude or temporal structure of the artificial DAN activation, thereby preventing the establishment of an appropriately quality-specific memory. Consistent with this possibility, whole-brain connectomic analyses have shown that larval DANs are among the most recurrently connected neurons in the brain (Winding et al 2023). Moreover, convergent one-step feedback from multiple MBONs innervating functionally distinct MB compartments is selectively routed onto individual larval DANs, suggesting that such recurrent architecture may contribute to the qualitative identity of the teaching signals (Eschbach et al 2020). In addition, it is important to emphasize that artificially induced memories have direct functional consequences for larval behavior. Identity-specific natural memories may support context-appropriate behavioral flexibility, allowing larvae to express avoidance only when the relevant aversive stimulus is present. By contrast, artificially imprinted memories may drive persistent and unconditional avoidance, leading larvae to reject contexts or food substrates that are otherwise harmless.

### Tonic baseline activity of DANs as a bidirectional teaching signal

A central finding of this study is that optogenetic silencing of DAN-f1/DAN-g1 during odor presentation was sufficient to induce appetitive memory (Figure 5D). This observation suggests that a decrease in tonic DAN activity can be interpreted by the MB circuit as a reward-like signal. In the larval MB, current models propose that dopaminergic DL1 and pPAM clusters are functionally balanced. Activation of punishment-encoding DL1 DANs is thought to perturb this balance and bias net MB output toward aversive valence, partly through inhibition of approach-promoting MBONs (Eschbach et al 2021, Eschbach & Zlatic 2020, Jurgensen et al 2024, Thum & Gerber 2019, Weber et al 2023a). Conversely, activation of reward-encoding pPAM DANs shifts MB output toward appetitive valence. A similar mechanism has been demonstrated in numerous studies of the adult MB, despite its substantially larger neuronal substrate, which comprises thousands rather than hundreds of neurons (Hige et al 2015, Owald et al 2015, Perisse et al 2016, Sejourne et al 2011).

Our results therefore suggest that, in addition to stimulus-dependent DAN activation, suppression of baseline DAN activity may represent an alternative mechanism for driving associative learning in the MB. If larval DL1 and pPAM clusters exhibit ongoing baseline activity that is mutually balanced under neutral environmental conditions, then selective silencing of punishment-encoding DANs in a rewarding context would be expected to disrupt this equilibrium and shift net MB output toward appetitive valence. Tonic dopaminergic signaling has been reported for several MB-projecting DAN classes in adult *Drosophila* exhibiting ongoing activity in the absence of acute reinforcement or sensory stimulation, including MP1, MV1 and PAM-γ3 DANs (Berry et al 2012, Placais et al 2012, Yamagata et al 2016). Even a similar mechanism has been reported in adult *Drosophila*; sugar reward suppresses baseline activity in PAM-γ3 dopaminergic neurons, thereby providing a reward signal during olfactory conditioning (Yamagata et al 2016). These results even suggest similarities to the vertebrate midbrain dopamine systems, where both increases and decreases in firing relative to baseline encode valence by the bidirectional activity (Danjo et al 2014, Tobler et al 2003). Thus, the same functional mechanism appears to operate across developmental stages of *Drosophila*, insects and maybe even more complex vertebrate systems. This would suggest that the bidirectional DAN activity function reflects a fundamental principle of learning circuit logic, rather than an emergent property arising from increased circuit complexity.

### Limitations of the computational model

The mechanistic model captures key behavioral phenomena using biologically grounded parameters derived from calcium imaging data, but several simplifications merit discussion. First, we assumed that KC–MBON plasticity is exclusively depressive, in line with dominant experimental findings (Hige et al 2015, Eschbach et al 2020). However, synaptic facilitation has been reported in other contexts (Cohn et al 2015, Handler et al 2019) and may contribute to valence reversal under different stimulus regimes without requiring cross-compartment interactions. Second, the model uses a single DAN and MBON per compartment, whereas the actual circuit contains multiple modulatory inputs and MBONs most compartments (Eichler et al 2017, Winding et al 2023). Additional modulatory neurons, dopaminergic or otherwise, could shape compartment-specific learning rules in ways not captured here. Third, allowing different learning rules in the individual compartments, in addition to innervation by different neurons, would also significantly alter the behavior of the model. Fourth, baseline DAN activities in the model are estimated from calcium imaging, which cannot resolve true resting firing rates, introduces slow indicator kinetics, and may misestimate the relative contributions of different compartments. Fifth, because calcium imaging was used to fit DAN neuron parameters, temporal dynamics on the sub-second timescale are not accurately represented; this is unlikely to affect minutes-long behavioral learning experiments but would need to be addressed for models of rapid online learning. Finally, the model does not simulate larval locomotion or the test-phase position dynamics of larvae navigating two odor sources-factors that are formally required to predict choice behavior from synaptic state and that represent a natural extension of the current framework using locomotion models (Jurgensen et al 2024, Sakagiannis et al 2026, Schleyer et al 2015).

The mechanistic MB model revealed that parallel, independent operation of individual DL1 compartments is insufficient to reproduce the full behavioral repertoire-in particular, valence reversal under combined salt punishment and DAN-f1/g1 silencing. Only model variants that incorporated cross-compartment interactions could account for this result. Specifically, Model 5 (Figure 10E), in which MBON output from the f1/g1 compartments provides excitatory feedback onto pPAM DANs, successfully replicated valence reversal. This cross-compartment motif is anatomically grounded in the larval connectome (Eichler et al 2017, Eschbach et al 2020). In addition, it conceptually aligns with evidence for compartmental crosstalk in adult *Drosophila* learning and extinction (Felsenberg et al 2018, Springer & Nawrot 2021) and second-order conditioning (Rachad et al 2025). The present data suggest that cross-compartment excitation of pPAM neurons by aversive-compartment MBON output is a functional circuit motif that can convert disrupted aversive signaling into active reward encoding. Importantly, this excitation must be tonically balanced by inhibitory input-presumably via c1/d1 MBON output onto pPAM-to prevent spurious appetitive memory in the absence of any manipulation. This balance-disruption mechanism offers a parsimonious account of how a single circuit can flexibly encode both aversive and appetitive memories depending on the configuration of DAN activity, without requiring changes in the learning rule itself.

## Methods

### Fly strains

Fly stocks were raised and maintained on standardized *Drosophila* food under controlled environmental conditions: 25°C, 60–80% humidity, and a 14/7-hour light/dark cycle (Weber et al 2023a, Weber et al 2023b). For behavioral analyses of DL1 DANs, the following split-GAL4 driver lines were used: MB054B (DAN-f1/DAN-g1, BDSC#603438), SS01716 (DAN-g1, BDSC#604149), SS02180 (DAN-f1, BDSC#603383), MB328B (DAN-d1, BDSC#602953), SS02160 (DAN-c1, BDSC#602922), and MB065B (DAN-c1, DAN-f1, BDSC#68281) (Eschbach et al 2020). To inhibit neuronal activity, split-GAL4 lines were crossed with the UAS-GtACR2 effector line (Mohammad et al 2017) (Bloomington stock center no. 92984), with experimental groups raised on standard *Drosophila* food supplemented with 0.5 mM all-*trans* retinal (ATR, Sigma Aldrich, cat. no. R2500). For neuronal activation experiments, split-GAL4 MB054B was crossed to UAS-ChR2XXL (Dawydow et al 2014) (Bloomington stock center no. 58374). Additionally, w¹¹¹⁸ flies (Bloomington stock center no. 3605) were used to obtain heterozygous effector controls. To ensure consistent experimental conditions, all genetic crosses were kept in darkness before the optogenetic experiments. Any handling of the larvae was performed under red light conditions.

### Gustatory choice behavior

Choice behavior assays were conducted as previously described (Hendel et al 2005, Huser et al 2012, Weber et al 2023a, Widmann et al 2016). Gustatory preference plates (85 mm diameter, Sarstedt, cat. no. 82.147) were prepared by dividing them into two halves: one containing a 2.5% (w/v) agarose solution (Sigma Aldrich, cat. no. A9539) and the other supplemented with either 1.5 M sodium chloride (VWR Chemicals, cat. no. 27810.364) or 2.0 M D-fructose (Sigma Aldrich cat. no. 47740) dissolved in 2.5% agarose. Plates were allowed to cool down at room temperature before testing. Approximately 30 third-instar larvae (feeding stage) were collected and placed at the center of the test plate, where they were allowed to move freely for 5 minutes.

For two-choice assays involving artificial blue light (470 nm) stimulation, preference plates were illuminated at either 220 lux to induce neuronal activation or 1100 lux to acutely block neuronal activity during the test, except for dark control groups. After the 5-minute testing period, larvae on each side of the plate were counted, and a gustatory preference index was calculated as follows:

Gustatory Preference Index:

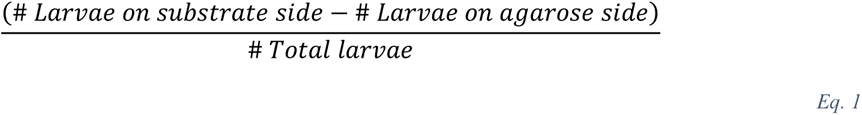

Positive scores indicate attraction, whereas negative scores indicate aversion.

### Olfactory choice behavior

Pure agarose (2.5% w/v) plates were cast as previously described (Hendel et al 2005, Huser et al 2012, Michels et al 2011, Weber et al 2023a, Weber et al 2023b, Widmann et al 2016). Custom made odor containers (Scherer et al 2003) filled with 10 µL AM (AM, Sigma Aldrich, cat. no. 46022, 1:250 diluted in paraffin oil (Sigma Aldrich, cat. no. 76235)) or 10 µL BA (undiluted, BA, Sigma Aldrich, cat. no. 12010) were placed on one side of a Petri dish plate (85 mm diameter, Sarstedt, cat. no. 82.1472) containing 2.5% (w/v) pure agarose. An empty container was placed on the opposite side to exclude visual or other side effects. Approximately 30 five to seven days old third instar feeding stage larvae were collected and placed on the test plate for 5 min. After this time, larvae were counted, subdivided into larvae on the odor side (#Odor) and larvae on the empty container side (#Empty). A preference index was calculated as follows:

Olfactory Preference Index:

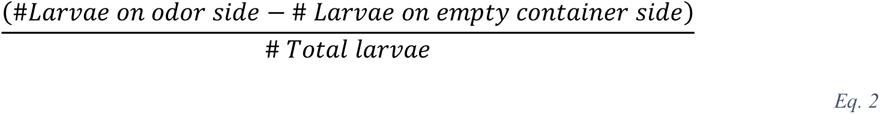

Positive scores indicate an appetitive preference, whereas negative scores indicate aversive preference.

### Olfactory learning and memory assays

Experiments were performed using standard methods (Eschbach et al 2020, Lyutova et al 2019, Schleyer et al 2020, Schroll et al 2006, Weber et al 2023a, Weber et al 2023b). Learning experiments were conducted on assay plates (85 mm diameter) filled with a thin layer of either 2.5% (w/v) pure agarose solution or 2.5% (w/v) agarose plus either 1.5 M sodium chloride solution, 2 M D-Fructose solution or 10 mM quinine solution (quinine hemisulfate; Sigma Aldrich cat. no. Q1250). Before closing the lids, solutions were allowed to cool down at room temperature to prevent condensation. As olfactory stimuli, we used amyl acetate (AM) diluted 1:250 in paraffin oil and undiluted benzaldehyde (BA). In both cases 10 μL odor were loaded into custom-made Teflon containers (4.5 mm diameter) with perforated lids (Scherer et al 2003).

A first group of approximately 30 third instar larvae (feeding stage) were collected and exposed to a first odor (AM+) while crawling on high-salt containing agarose for five minutes, followed by five minutes exposition to the second odor (BA-) on pure agarose medium. A second group of larvae received the reciprocal training paradigm (AM-/BA+). After one training cycle, larvae were transferred onto test plates containing agarose supplemented with high-salt on which odor AM and odor BA were presented on opposite sides. After five minutes, larvae were counted as located on the AM side (#AM), the BA side (#BA) or in a central portion of the dish (∼10mm wide neutral zone). Preference indices (PREF) for each training group were calculated as follows:

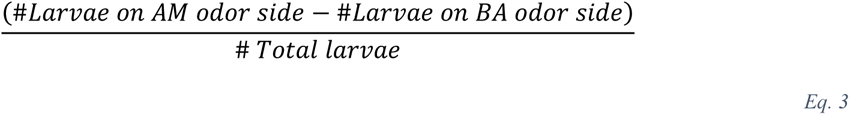

PREF (AM-/BA+):

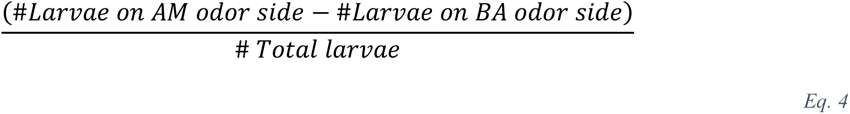

To measure specifically the effect of associative learning, a performance index (PI) was calculated as the difference in preference between reciprocally trained larvae:

Performance Index (PI):

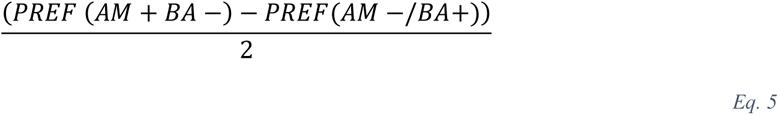

Negative scores indicate an aversive associated learning. By the division of two, scores were bound within -1;1. In addition we altered the sequence of odor presentation across repetitions of experiments.

### Optogenetic activation assays

To investigate the effects of artificial activation of DAN-f1 and DAN-g1 during different phases of the two-odor high-salt learning paradigm, the split-GAL4 line MB054B was crossed with the effector line UAS-ChR2XXL. Flies were tested under two conditions: in darkness (dark control) or with blue light activation (470 nm, 220 lux, experimental group). Blue light stimulation was applied at specific phases: (i) during the entire training phase of five minutes on the pure agarose plate, (ii) during the entire training phase of five minutes on the high-salt plate, (iii) throughout the entire training that consists of the two five minute phases mentioned before, and (iv) during the entire test phase of five minutes on the high-salt plate. Additionally, w1118 flies were crossed with UAS-ChR2XXL to generate a heterozygous effector control group, which underwent the same experimental conditions as the experimental group.

To evaluate whether artificial blue light activation can substitute for an aversive stimulus, the same genetic crossings were used. However, larvae were subjected to a modified learning paradigm. Here, the negative gustatory reinforcer was replaced with a pure agarose plate, and blue light stimulation was applied instead. Additionally, larvae were tested on different substrates containing either pure agarose (2.5%, w/v), agarose supplemented with 10 mM quinine, or agarose supplemented with 2.0 M D-fructose. Preference and performance indices were calculated as described previously.

### Optogenetic inhibition experiments

To acutely block synaptic output, we used UAS-GtACR2. We crossed the effector to the DL1 specific split-GAL4 driver lines as mentioned before. Flies were maintained on standard food supplemented with 0.5 mM all-*trans*-retinal at 25°C as described before (Meloni et al 2020). Vials were wrapped in aluminum foil to ensure larval development in constant darkness. Groups of about 30 third instar feeding stage larvae received a reciprocal two-odor training regime as above. Larvae were exposed to blue light (470 nm, 1100 lux) during different phases throughout the behavioral paradigm: (i) during the entire training phase of five minutes on the pure agarose plate, (ii) during the entire training phase of five minutes on the high-salt plate, (iii) throughout the entire training that consists of the two five minute phases mentioned before, and (iv) during the entire test phase of five minutes on the high-salt plate.

In one experimental condition, larvae were subjected to a modified training paradigm. They were trained exclusively on plain agarose plates, with blue light stimulation delivered during only one of the two 5-minute training phases, so that neuronal inhibition occurred in conjunction with one odor presentation but not the other. Subsequent testing was performed on a high-salt containing plate under dark conditions. Control experiments were conducted using the same genotype but with standard food lacking 0.5 mM all-*trans* retinal. The data were then calculated and scored as mentioned above.

### Statistical analysis

Results were analyzed using GraphPad Prism 8.4.3. Datasets were assessed for normal distribution by using the Shapiro-Wilk test. Parametric Datasets were further analyzed by using either unpaired t-test (comparison of two groups) or one-way ANOVA (comparison of more than two groups). One sample t-tests were performed to compare medians against chance level of parametric groups. Non-parametric datasets were tested with Mann-Whitney-U test (comparison of two groups) or Kruskal-Wallis followed by Dunn’s multiple comparison (comparison of groups larger than two). Moreover, Wilcoxon sign-ranked test was performed to test medians of non-parametric groups against chance level.

Results are visualized as box-plots, indicating the median as middle line, 25%/75% quantiles as box boundaries and minimum/maximum performance indices as whiskers. Each data point is represented as black dot and sample sizes are noted within graphics. Asterisks and ‘‘n.s.’’ indicate p < 0.05 and p > 0.05, respectively. The source data and results of all statistical tests are documented in the Supplemental Information file.

### Simulation of the computational model

In our network model of the larval MB (Figure 9A), a population of 100 KCs encode the odor stimulus and make excitatory, plastic connections with MBONs. The synaptic weights *wij* onto each MBON are subject to modulation by a single DAN (Figure 9), reflecting the compartmentalized organization of the MB lobes (Aso et al 2014, Eichler et al 2017, Hige et al 2015). In the insect MB, KCs show very low spontaneous activity in experimental recordings (Turner et al 2008) and respond to odors in a population-sparse manner (Honegger et al 2011, Ito et al 2008, Perez-Orive et al 2002, Turner et al 2008). Accordingly, between 5% and 10% of the model KC population is activated by any given odor, with individual activation levels drawn from a uniform distribution for each KC (Eq. 6, Figure 9B, Table 1), following the approach in earlier models (Manoim Wolkovitz et al 2026). KC responses to a virtual odor stimulation are modeled as time-varying activation functions that decay exponentially during the duration of the odor stimulation *T*stim from *t*startto *t*stop (Eq. 6) towards a steady state calcium activation *C*oo (Eq. 6). This is consistent with experimental KC recordings in insects (Bilz et al 2020, Demmer & Kloppenburg 2009, Froese et al 2014, Ito et al 2008, Szyszka et al 2005, Turner et al 2008) and KC responses in full olfactory pathway models in the larval (Jürgensen et al 2021, Jurgensen et al 2024) and adult fruit fly (Betkiewicz et al 2020, Rapp & Nawrot 2020). For each odor-activated KC in the model, an initial activation level is drawn from a uniform distribution (Eq. 6, Table 1) and onset and offset of KC odor responses occur with random delays, likewise drawn from uniform distributions (Table 1, Eq. 6). Each DAN (c1, d1, f1, g1, pPAM, Eq. 7) and MBON (Eq. 8) is represented by a differential equation that describes the time-varying calcium activation level *c* in arbitrary units as determined by the integration of synaptic inputs. Adaptation is captured in the DAN neuron models in the adaptation term *a* (Eq. 7). The four DL1-DANs are modelled as individual neurons (c1, d1, f1, g1), while the pPAM neurons are collectively modelled as one single neuron, collapsing the entire cluster of four neurons into one single model neuron, due to availability of recordings (Weber et al., 2025). Each approach-mediating MBON+ represents the output of a single MB compartment, innervated by one respective DL1-DAN (c1, d1, f1, g1, Figure 9A). The avoidance-mediating MBONs are innervated by the neurons of the pPAM cluster. The implemented connectivity between KCs and each MBON is dense; each postsynaptic MBON *i* is connected with all KCs *j*, but only odor-activated presynaptic KCs deliver input into the respective MBON as there is no baseline KC activity. The KC-MBON synapses in each compartment (c1, d1, f1, g1, pPAM) are innervated by the respective single DAN or pPAM cluster. The coincidence of odor-driven KC activity and DAN activity results in the modulation of the synaptic weight *wij* of the respective KCj-MBONi synapse, proportional to the learning rate *lr* (Eq. 9), with only depression permitted. The MBONs project onto descending pathways that control animal behavior and are usually conceptualized as approach- or avoidance-mediating (Aso et al 2014, Eschbach et al 2021, Owald et al 2015, Saumweber et al 2018, Sejourne et al 2011, Thum & Gerber 2019).

**Table 1:** Neuron and network model parameters.

| Parameter | Value | Variable name |
| --- | --- | --- |
| dt | 1sec | dt |
| Number of KCs | 100 | n_KC |
| Number of MBON<sup>+</sup> | 4 | n_MBONp |
| Number of MBON<sup>-</sup> | 1 | n_MBONn |
| Initial weight KC-MBON<sup>+</sup> | 0.7 | w_KC_MBONp |
| Initial weight KC-MBON<sup>+</sup> | 0.7*4 | w_KC_MBONn |
| MBON time constant | 0.5sec | tau_MBON |
| pPAM DAN time constant | 0.637312sec | tau_DAN_PAM |
| pPAM adaptation factor | 0.193716 | b_PAM |
| pPAM adaptation time constant | 3.243857sec | tau_a_DAN_PAM |
| c1 time constant | 8.952809sec | tau_DAN_c1 |
| c1 adaptation factor | 13.75433 | b_c1 |
| c1 adaptation time constant | 0.865808sec | tau_a_DAN_c1 |
| d1 time constant | 11.550999sec | tau_DAN_d1 |
| d1 adaptation factor | 10.237083 | b_d1 |
| d1 adaptation time constant | 1.24174sec | tau_a_DAN_d1 |
| f1 time constant | 0.579765sec | tau_DAN_f1 |
| f1 adaptation factor | 0.006707 | b_f1 |
| f1 adaptation time constant | 61.637608sec | tau_a_DAN_f1 |
| g1 time constant | 0.667098sec | tau_DAN_g1 |
| g1 adaptation factor | 0.273706 | b_g1 |
| g1 adaptation time constant | 32.700818sec | tau_a_DAN_g1 |
| Input weight salt c1 | 0.826388 | w_pun_DANc1 |
| Input weight salt d1 | 1.002981 | w_pun_DANd1 |
| Input weight salt f1 | 0.012125 | w_pun_DANf1 |
| Input weight salt g1 | 0.061857 | w_pun_DANg1 |
| Input weight salt pPAM | -0.175809 | w_pun_DANPAM |
| Input weight sugar c1 | 0.54274 | w_rew_DANc1 |
| Input weight sugar d1 | 0.093441 | w_rew_DANd1 |
| Input weight sugar f1 | -0.023435 | w_rew_DANf1 |
| Input weight sugar g1 | -0.011659 | w_rew_DANg1 |
| Input weight sugar pPAM | 0.379457 | w_rew_DANPAM |
| **Variable parameter** |  |  |
| Learning rate | $lr \sim U(-0.0003, -0.0001)$ | lr |
| DAN baseline activity | $bl \sim U(0.01, 0.03)$ | bl |
| Percentage odor activated KCs | $pc\_active\_KC \sim U(5, 10)$ | pc_active_KC |
| KC odor response | $activation \sim U(0.1, 0.15)$ | activation |
| Odor response onset | $start\_offset \sim U(0sec, 5sec)$ | start_offset |
| Odor response offset | stop_offset $\sim U(-5\text{sec}, 5\text{sec})$ | stop_offset |
| Exponential decay constant | 9.00 | $\lambda$ |
| Steady state activity | 0.06 | $C_{\infty}$ |
| **Additional parameters models 2-5** |  |  |
| Weight MBON-MBON (M3) | w_MBON_MBON $\sim U(-0.1, -0.01)$ | w_MBON_MBON |
| Weight MBON-DAN (M4) | w_MBON_DAN $\sim U(0.1, 1.0)$ | w_MBON_DAN |
| Weight MBONn-pPAM (M4) |  | -w_MBON_DAN |
| Weight MBONp-DI1 (M4) |  | -w_MBON_DAN |
| Weight MBONn-DL1 (M4) |  | w_MBON_DAN/4 |
| Weight MBONp-DAN (M5) | w_MBONp_DAN $\sim U(5.0, 8.0)$ | w_MBONp_DAN |

In our model, all MBON activity is integrated by the descending pathway for the control of avoidance behavior. It receives excitatory input from the avoidance-mediating MBON- and is inhibited by approach-mediating MBONs+. The avoidance bias is the time-averaged difference between the total activity of MBON- and MBONs+, as integrated across the test phase from *t*startto *t*stop (Eq. 10). Positive values of the avoidance bias indicate a model-predicted shift toward odor avoidance, whereas negative values indicate a shift toward odor approach.

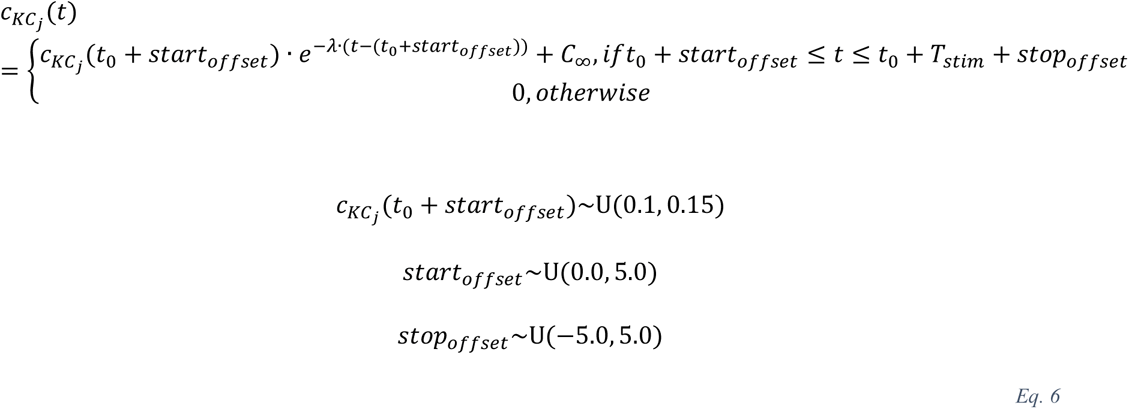

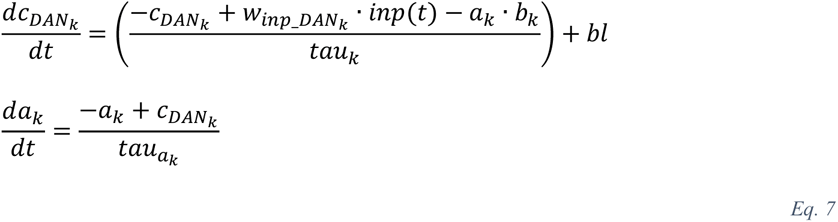

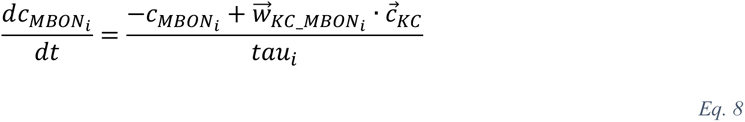

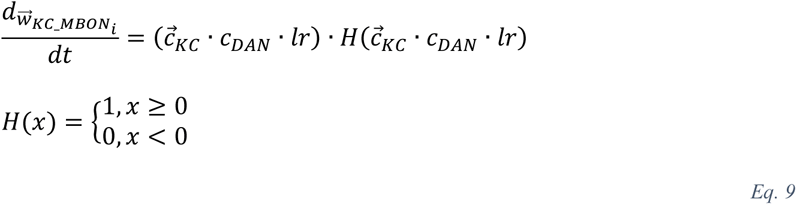

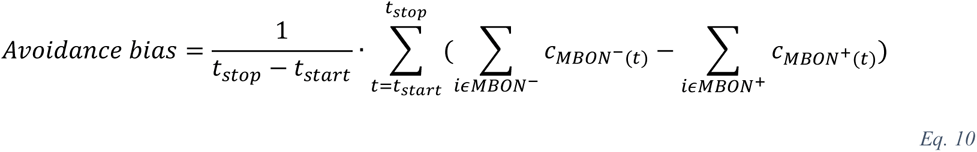

Simulations of each experiment involved 200 model instances with some parameters randomly drawn from uniform distributions to assess the model’s robustness (Table 1). Each experiment lasted six minutes, simulated with a time resolution of one second. Odor stimuli activated the KCs for 5 minutes. In addition to odor-driven activation (i) DANs could be activated by a high-salt stimulus or through simulated optogenetic activation (DAN-f1 and g1), or (ii) DANs could be silenced (DAN-f1 and g1). Salt-induced DAN activation occurred via binary synaptic input denoting reinforcement on or off, respectively. Optogenetic activation and silencing were implemented as additional excitatory or inhibitory synaptic inputs to the DANs.

In addition to the base model (Figure 9A), four extended versions of the base model were implemented (Figure 10-figure supplement 1). We maintained all neurons, connections and parameters present in the base model. In model 2 (Figure 10-figure supplement 1A), DAN-f1 is not activated by high-salt and thus does not contribute to reinforcement-driven plasticity in the MB. It does, however, exhibit spontaneous activity. In model 3 (Figure 10figure supplement 1B), there are additional inhibitory synapses between MBONs of approach and avoidance compartments. Each MBON+ receives inhibitory input from each MBON- and vice versa. These interactions can be interpreted as either direct or indirect, via interneurons for net inhibitory effects of MBONs. Model 4 (Figure 10-figure supplement 1C) features inhibitory MBON-DAN feedback is added within compartment, and excitatory between compartments. The pPAM cluster receives inhibitory synaptic input from each MBON- and excitatory input from each MBON+. Each DL1-DAN receives excitatory synaptic input from the population of MBONs- and inhibitory within compartment input from the respective MBON+. Model 5 (Figure 10-figure supplement 1D) includes cross-compartment MBON+-pPAM feedback. MBONs of the c1 and d1 compartment inhibit the pPAM cluster, while the feedback from the MBONs of the f1 and g1 compartment is excitatory.

### Fitting of neuron model parameters to experimental data

To obtain biologically realistic parameters for each DAN (DL1-DANs c1, d1, f1, g1 and the pPAM cluster) we independently fitted the neuron model parameters (*tau, b, taua, winp_DAN*, Table 1) to match the time-resolved experimental calcium-imaging traces (Figure 8) averaged across animals taken from (Weber et al 2025). Reinforcement input into a DAN was encoded as time-resolved binary input (on/off) as the experimentally applied concentration is assumed to be constant (Weber et al 2025). For the DL1-DANs, we first used the high-salt response calcium imaging data to fit all parameters of the respective neuron model (*tau, b, taua, winp_DAN*) and then used the fructose-response calcium-imaging data to fit the input weight *winp_DAN* again for the response to fructose with all other parameters (*tau, b, taua*) kept constant. For the pPAM cluster, represented by a single DAN in our model (Figure 9A), we conversely performed a parallel fit of all parameters (*tau, b, taua, winp_DAN*) on the experimental calcium-imaging of the PAM cluster to fructose stimulation, and in a second step fitted the input weight based on the high-salt response. Parameters were estimated by minimizing the sum of squared residuals between the model output and the observed calcium trace with length *m*.

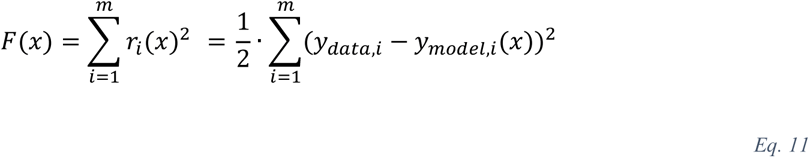

Initial values for each parameter were randomly drawn from a uniform distribution from a minimum to a maximum initial value (*tau* [0.5, inf], *b* [0, inf], *taua* [0.5, inf], *winp_DAN* [0, 10]). We performed 10,000 optimizations, each with a randomly drawn set of initial parameters and selected the resulting parameter configuration with the smallest sum of squared error (Table 1). Then the input weight of the DL1-DANs were fitted to match the fructose response data and salt response for the pPAM cluster. The initial *w* was again selected from a uniform distribution (*winp_DAN* [-20, 20]).

## Data availability statement and Supplemental Information

Supplemental Information includes supplemental experimental procedures, statistical evaluations, and additional figures and can be found with this article online.

The code for the model implementation can be obtained at https://github.com/nawrotlab/distributed_DAN_signal

## Acknowledgements

This work was supported by the Deutsche Forschungsgemeinschaft (grant no. 441181781, 426722269, 432195391 to AST and 365082554 to MPN), by EU funds from the ESF Plus Program (Grant No. 100649752) all to AST and by the Federal Ministry of Education and Research (BMBF, grant no. 01GQ2103A) to MPN. AMJ received additional funding from the European Union’s Horizon research and innovation programme under the Marie Skłodowska-Curie (grant agreement No. 101205002). We thank Bert Klagges, Tilman Triphan, Dennis Pauls, and Wolf Huetteroth for discussions and comments. Additionally, we thank Astrid Rohwedder for fly care and maintenance.

## Author contributions

Conceptualization, D.W., A-M.J., M.P.N., and A.S.T; Methodology, D.W., J.K., A-M.J., M.P.N., and A.S.T; Investigation, D.W., J.K., and A-M.J.; Writing, D.W., A-M.J., M.P.N., and A.S.T; Supervision, D.W., A-M.J., M.P.N., and A.S.T.

---

**Figure 7- Figure supplement 1:**
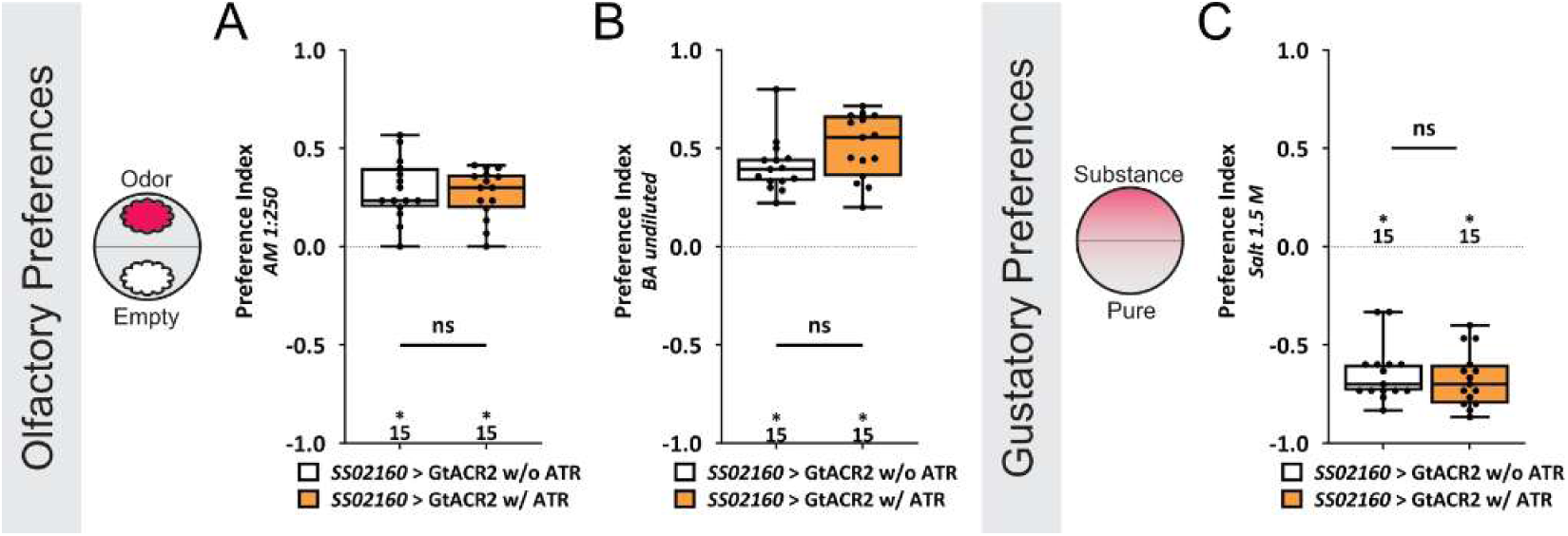
Acute optogenetic inhibition of DAN-c1 does not change the naïve chemosensory responses of *Drosophila* larvae. To determine whether optogenetic inactivation of DL1 DAN-c1 changes the chemosensory choice behavior of naïve larvae, the SS02160 split-GAL4 driver together with UAS-GtACR2 was used. For all experiments two groups were tested: SS02160 split-GAL4; UAS-GtACR2 control larvae that did not receive all-*trans* retinal in their diet (white) and SS02160 split-GAL4; UAS-GtACR2 experimental larvae that were fed all-*trans* retinal throughout development (orange). (A-C) In all three cases the experimental animals showed a clear olfactory attraction (A,B; p < 0.05) or gustatory avoidance (C; p < 0.05), which is comparable to the respective control group (p > 0.05). All behavioral data is shown as box-plots. Differences between groups are highlighted by horizontal lines between them. Performance indices different from random distribution are indicated below (in A and B) or above (in C) each box-plot. The sample size of each group (N=15) is given for each box-plot. n.s. p > 0.05; * p < 0.05. The source data and results of all statistical tests are documented in Figure 7 Figure supplement 1 — source data 1.

**Figure 10- Figure supplement 1:**
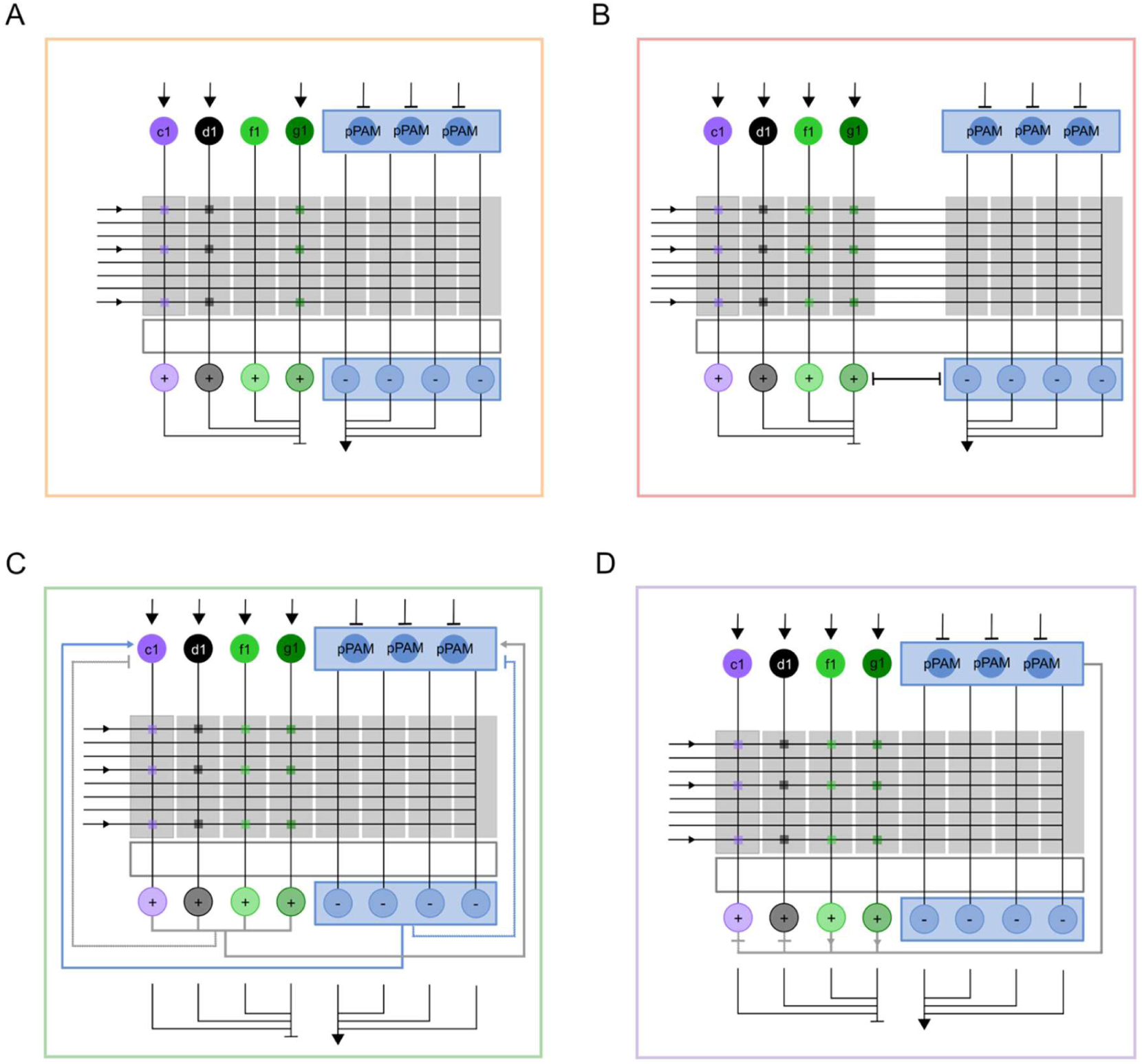
All extended circuit models. All models are based on base model 1 (Figure 9A). (A) In model 2, salt-driven activation of DAN-f1 is omitted. (B) Model 3 features bidirectional inhibitory connections between approach- and avoidance-promoting MBONs. (C) Model 4 includes additional MBON-DAN feedback within compartment, and excitatory between compartments. (D) And in model 5, the pPAM cluster received an mix of inhibitory and excitatory inputs from different MBONs+ (MBONs of the c1 and d1 compartment inhibit the pPAM cluster, while the feedback from the MBONs of the f1 and g1 compartment is excitatory).

